# Neocentromere formation through Robertsonian fusion and centromere repositioning during the evolution of zebras

**DOI:** 10.1101/2022.02.15.480582

**Authors:** Eleonora Cappelletti, Francesca M. Piras, Lorenzo Sola, Marco Santagostino, Wasma A. Abdelgadir, Elena Raimondi, Solomon G. Nergadze, Elena Giulotto

## Abstract

Centromeres are epigenetically specified by the histone H3 variant CENP-A and typically associated to highly repetitive satellite DNA. We previously discovered natural satellite-free neocentromeres in *Equus caballus* and *E. asinus*. Here, through ChIP-seq with an anti-CENP-A antibody, we found an extraordinarily high number of centromeres lacking satellite DNA in the zebras *E. burchelli* (15 of 22) and *E. grevyi* (13 of 23), demonstrating that the absence of satellite DNA at the majority of centromeres is compatible with genome stability and species survival and challenging the role of satellite DNA in centromere function. Nine neocenstromeres are shared between the two species in agreement with their recent separation. We *de novo* assembled all neocentromeric regions and improved the reference genome of *E. burchelli*. Sequence analysis of the CENP-A binding domains revealed that they are LINE-1 and AT-rich with four of them showing DNA amplification. In the two zebras, satellite-free centromeres emerged from centromere repositioning or following Robertsonian fusion. In five chromosomes, the centromeric function arose near the fusion points, which are located within regions marked by traces of ancestral pericentromeric sequences. Therefore, besides centromere repositioning, Robertsonian fusions are an important source of satellite-free centromeres during evolution. Finally, in one case, a neocentromere was seeded on an inversion breakpoint. At eleven chromosomes, whose primary constrictions seemed to be associated to satellite repeats by cytogenetic analysis, neocentromeres were instead located near the ancestral inactivated satellite-based centromeres, therefore, the centromeric function has shifted away from a satellite repeat containing locus to a satellite-free new position.

## INTRODUCTION

Centromeres are essential nucleoprotein structures of eukaryotic chromosomes responsible for the correct segregation of sister chromatids during cell division. Centromeres are epigenetically specified by the histone H3 variant CENP-A, the hallmark of a functional centromere (Allshire and Karpen 2008). Mammalian centromeres are typically associated to extended arrays of tandemly iterated sequences (satellite DNA), which are divergent and represent the most rapidly evolving components of genomes (Plohl, et al. 2014). The presence of such sequences has hampered comprehensive molecular analysis of these intriguing loci. Our discovery that equids are characterized by centromeres completely devoid of satellite DNA made these species an exceptional model system for dissecting the molecular architecture of mammalian centromeres as well as understanding the mechanisms driving centromere birth and maturation during evolution (Wade, et al. 2009; Piras, et al. 2010; Purgato, et al. 2015; Giulotto, et al. 2017; Nergadze, et al. 2018; Roberti, et al. 2019).

The family of Equidae, with its only extant genus, the *Equus* (horses, asses and zebras), belongs to the order Perissodactyla together with Tapiridae (tapirs) and Rhinocerotidae (rhinoceroses). While the karyotypes of Tapiridae and Rhinocerotidae remained quite stable during evolution and resemble the putative perissodactyl ancestral karyotype, characterized by high chromosomal number and a prevalence of acrocentric chromosomes (Trifonov, et al. 2008), *Equus* karyotypes underwent a rapid evolution after their divergence from the common ancestor, dated around 4 million years ago (Mya) (Orlando, et al. 2013). The most recent radiation events, which differentiated asses and zebra species, date back to less than 1 Mya and many species and subspecies emerged in this short evolutionary time (Trifonov, et al. 2008; Trifonov, et al. 2012; Jónsson, et al. 2014). The extensive karyotype reshuffling that occurred during equids speciation is due to chromosome rearrangement and centromere repositioning, that is the shift of the centromeric function without sequence rearrangement (Carbone, et al. 2006; Piras, et al. 2010; Trifonov, et al. 2012). These phenomena led to the evolution from the ancestral karyotype, with the majority of chromosomes being acrocentric, to karyotypes with a reduced chromosomal number, mainly composed by meta- and sub-metacentrics. Our previous cytogenetic analyses suggested that several of these chromosomes harbor satellite-free centromeres (Carbone, et al. 2006; Piras, et al. 2010) while displaying blocks of satellite DNA sequences at non-centromeric chromosome ends, as relics of the inactivated centromeres from the ancestral acrocentrics. We then characterized, at molecular level, neocentromeres deriving from centromere repositioning events in horse and donkey (Wade, et al. 2009; Nergadze, et al. 2018). The position of these peculiar centromeres, identified as CENP-A binding domains, can slide within a relatively wide region, probably limited by epigenetic boundaries. These domains were defined epialleles and the phenomenon was called centromere sliding (Purgato, et al. 2015; Nergadze, et al. 2018).

ChIP-seq experiments performed with an anti-CENP-A antibody on individuals from families composed by donkeys, horses and their hybrid offspring (mule/hinny) revealed that such epialleles are inherited as Mendelian traits, but their position can slide in one generation (Wade, et al. 2009; Nergadze, et al. 2018). Conversely, the position of the centromere is stable during mitotic propagation of cultured cells grown for several population doublings, suggesting that the sliding can presumably take place during meiosis (Wade, et al. 2009; Nergadze, et al. 2018).

Sequence analysis of the donkey satellite-free centromeric domains demonstrated that five neocentromeres were characterized by novel tandem repetitions. These amplified genomic sequences are chromosome specific, with amplified units ranging in size from a few to tens of kilobases (Wade, et al. 2009; Nergadze, et al. 2018). The repeat copy number was variable in different individuals, suggesting the existence of polymorphism in the population. We suggested that these amplified DNA may represent an intermediate evolutionary stage toward satellite DNA formation during the process of centromere maturation (Wade, et al. 2009; Nergadze, et al. 2018). According to this model, after centromere inactivation, associated satellite sequences are maintained at the original site while the newly born centromere is completely devoid of satellite sequences. Subsequently, satellite DNA is gradually lost at the locus of the ancestral centromere, while, at the functional satellite-free centromere, sequence amplification may occur before complete maturation via satellite DNA acquisition.

Besides centromere repositioning, it is well described that Robertsonian translocations, which involve centric fusion of two acrocentric chromosomes, marked the rapid karyotype reshaping of zebras (Piras, et al. 2010; Musilova, et al. 2013), whose centromeres still lack a molecular characterization. According to current taxonomy, there are three zebra species, namely plains zebra (*Equus quagga*, also known as *Equus burchelli*), Grevy’s zebra (*Equus grevyi*) and mountain zebra (*Equus zebra*) (Ransom and Kaczensky 2016). In previous work, we described the localization of satellite DNA families by FISH in Burchell’s zebra and Grevy’s zebra, showing that several centromeres are devoid of satellite DNA at cytogenetic resolution (Piras, et al. 2010). The aim of the present work was to identify and characterize at molecular level satellite-free centromeres in the two zebras and to test whether Robertsonian fusions played any role in neocentromere formation.

## RESULTS

### Burchell’s zebra reference genome

In 2020, a *de novo* chromosome-length assembly for the Burchell’s zebra (Equus_quagga_HiC) was released by the DNA Zoo team (https://www.dnazoo.org). Since this assembly contains scaffolds corresponding to entire zebra chromosomes but lacking chromosome assignment, we performed a whole-genome alignment of the Equus_quagga_HiC assembly to the horse genome and assigned the scaffolds to specific horse chromosomes using EquCab3.0 as reference. Since the horse and the Burchell’s zebra genomes share high sequence identity (Jónsson, et al. 2014) and chromosome orthologies are well described (Musilova, et al. 2013), taking advantage of the sub-chromosomal comparative maps between horse and Burchell’s zebra (Musilova, et al. 2013), we assigned chromosome numbers to the 22 Burchell’s zebra chromosomal scaffolds (Supplementary Figure S1). The results showed that fourteen scaffolds were correctly oriented compared to the direction previously determined by comparative cytogenetics (Musilova, et al. 2013) whereas nine scaffolds, corresponding to chromosomes 2, 6, 9, 12, 13, 14, 16, 21 and X, had a reverse orientation.

### Identification and sequence assembly of satellite-free centromeres in Burchell’s zebra

In previous work we described several centromeres from *Equus burchelli* (Burchell’s zebra) lacking detectable satellite repeats at the cytogenetic level (Piras, et al. 2010). To identify satellite-free CENP-A binding domains of Burchell’s zebra at molecular level, a ChIP-seq experiment with an antibody against CENP-A was carried out on primary skin fibroblasts. The ChIP-seq reads were mapped both on the Burchell’s zebra (Equus_quagga_HiC) assembly and on the horse (EquCab3.0) reference genome, which remains the best assembled genome sequence among equids (Wade, et al. 2009; Kalbfleisch, et al. 2018). Using the pipeline that we developed for satellite-free centromeres in the donkey (Nergadze, et al. 2018), genomic regions, enriched for CENP-A binding, were identified. In Figure 1A and B, the graphical representation of the CENP-A enrichment peaks found using as reference EquCab3.0 or Equus_quagga_HiC, respectively, is shown. Map positions of the enrichment peaks are reported in Supplementary Table S1. These peaks, which correspond to satellite-free centromeres, were located on 15 out of the 22 chromosomes. Thirteen of these chromosomes (4, 6, 7, 8, 9, 10, 12, 14, 15, 16, 17, 18 and X) are meta- or sub-metacentric while two of them (20 and 21) are acrocentric.

**Figure 1.**
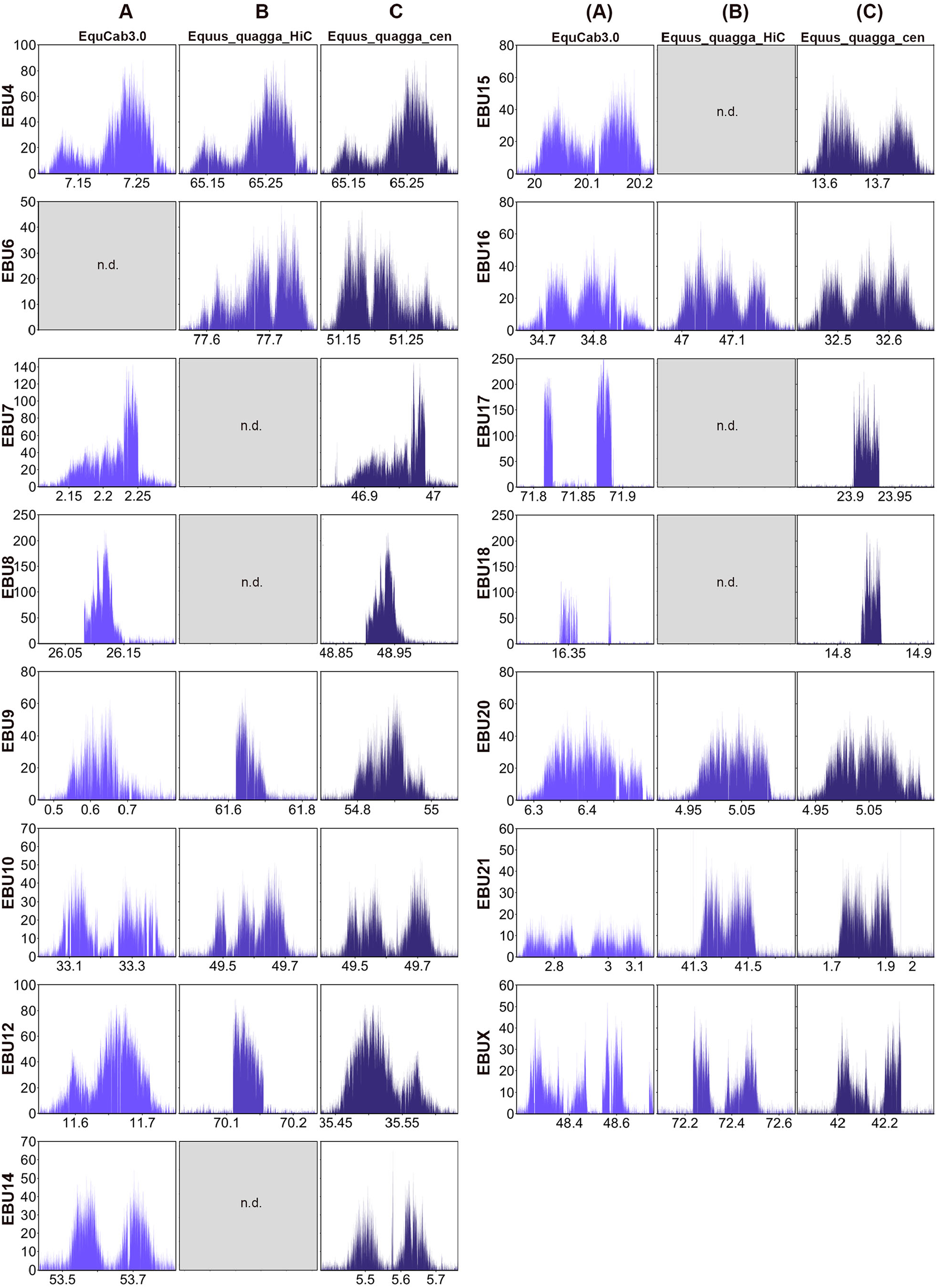
Satellite-free centromeres in Burchell’s zebra. ChIP-seq reads from primary fibroblasts of Burchell’s zebra were mapped on EquCab3.0 (A), Equus_quagga_HiC (B) and Equus_quagga_cen (C). CENP-A enriched domains are visualized as peaks. The y-axis reports the normalized read count whereas the x-axis reports the coordinates on the reference genome. For each chromosome, genomic windows of the same length are plotted.

Satellite-free CENP-A binding domains on chromosomes 4, 9, 10, 12, 16, 20, 21 and X were identified on orthologous positions in both reference genomes (Figure 1A and B, Supplementary Table S1). For chromosome 6, a CENP-A binding domain was detected only in the zebra assembly suggesting that the orthologous horse locus is highly rearranged. A detailed analysis of this centromere, which is reported in a following paragraph, confirms this hypothesis. On the contrary, the centromeric domains of chromosomes 7, 8, 14, 15, 17 and 18 were detected only in the horse reference genome suggesting that these loci are not properly assembled in the DNA Zoo Equus_quagga_HiC scaffolds.

Although some peaks showed a fairly regular Gaussian-like shape (such as those on chromosomes 4 and 20), some of them were irregular and contained several gaps on both reference genomes (such as those on chromosome 10 and 18). In addition, for several centromeres (such as 9 and 12), the shape and size of the peak were different between the two assemblies, reflecting sequence variation between the two species and/or sequence gaps.

To determine more precisely the organization of the CENP-A binding domains at satellite-free centromeres, the actual DNA sequence corresponding to the 15 Burchell’s zebra centromeres was assembled from our Illumina reads (see Materials and Methods). For each centromeric region, genomic segments ranging in size between 67 and 445 kb, containing the CENP-A binding domain, were obtained (accession numbers: OM643400-OM643414). We then corrected the Equus_quagga_HiC sequence by replacing the incomplete or misassembled centromeric loci with the newly assembled genomic segments of Burchell’s zebra. In addition, we adjusted the direction of the chromosomes which were incorrectly oriented in the DNA Zoo assembly (2, 6, 9, 12, 13, 14, 16, 21 and X) (Figure 2, Supplementary Table S2). Two of the newly oriented chromosomes (2 and 13) contain satellite-based centromeres. The resulting genome sequence is, from now on, called Equus_quagga_cen. In Figure 2, the pairwise genome comparison between EquCab3.0 and Equus_quagga_cen chromosomes is shown. The ChIP-seq reads from Burchell’s zebra were then mapped on this new reference genome and enrichment peaks, corresponding to CENP-A binding domains, were obtained (Figure 1C, Supplementary Table S3). The peak profiles visualized on the new reference genome showed that several gaps and irregular shapes were no longer detected and their profiles were improved (compare Figure 1 A, B and C). It is worth noticing that, when Equus_quagga_cen was used as reference, some CENP-A binding peaks (6, 12, 16 and X) showed a specular shape compared to the peaks obtained with the horse and Burchell’s zebra reference genomes. This effect is a consequence of the opposite orientation of these chromosomes in the Equus_quagga_cen assembly compared to the other reference genomes. In a few cases, such as EBU10, EBU14 and EBU15, two well separated or partially overlapping CENP-A binding domains are visible. They probably correspond to different epialleles on the two homologous chromosomes in the individual analyzed here and suggest positional variation of satellite-free centromeres in the population.

**Figure 2.**
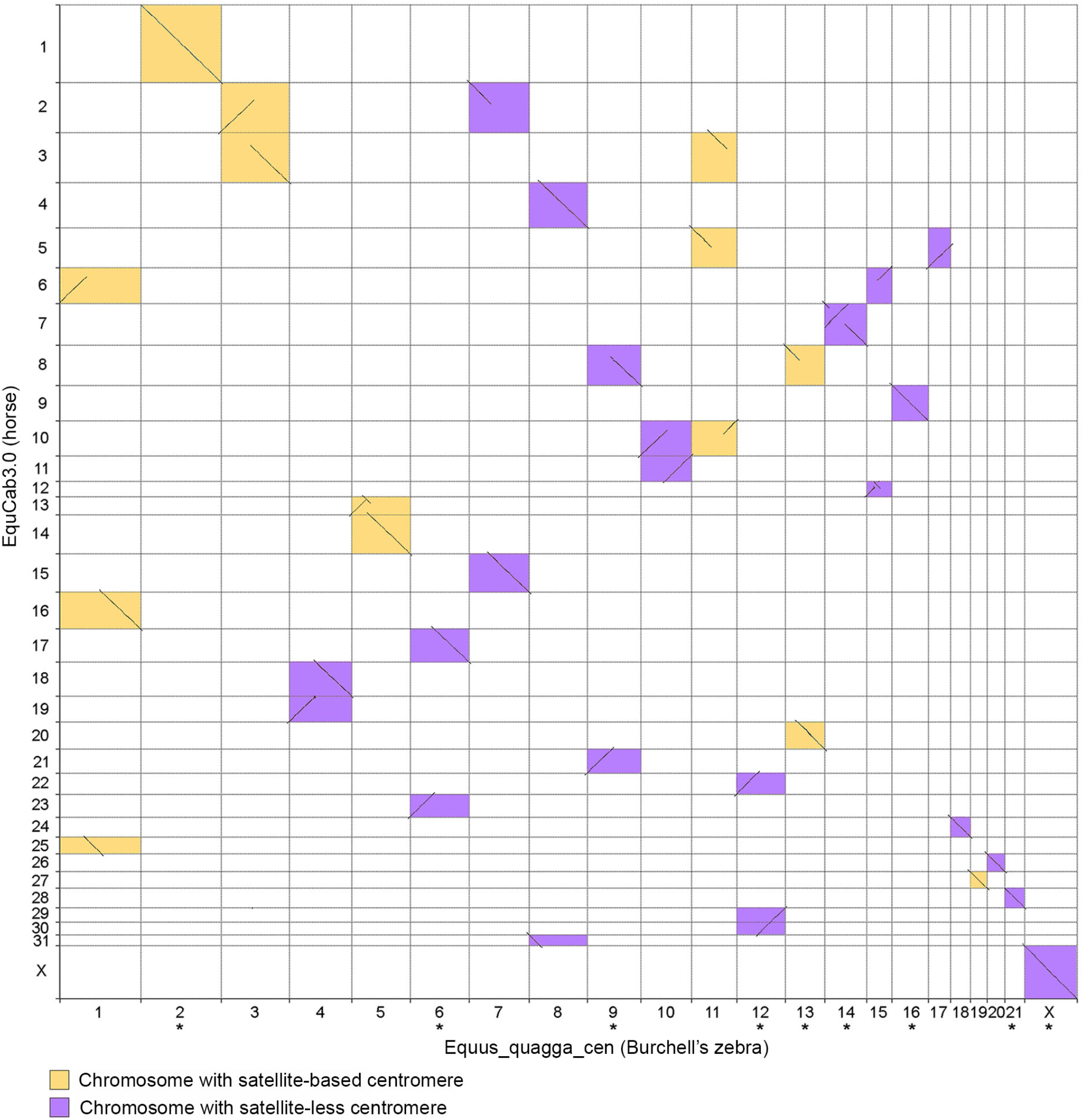
Pairwise genome comparison between EquCab3.0 and Equus_quagga_cen. Chromosome numbers were assigned to Equus_quagga_HiC scaffolds corresponding to entire zebra chromosomes. Aligned segments between horse (y-axis) and Burchell’s zebra (x-axis) chromosomes are represented as lines. Colors indicate chromosomes with satellite-free (purple) and satellite-based (yellow) centromeres. The orientation of the chromosomes marked with an asterisk was corrected according to the direction previously determined by cytogenetic analysis (Musilova, et al. 2013).

### Identification and sequence assembly of satellite-free centromeres in Grevy’s zebra

To describe at the molecular level Grevy’s zebra satellite-free centromeres, a ChIP-seq experiment was carried out using the anti-CENP-A antibody with Grevy’s zebra fibroblasts. Since a reference genome for this species is not available, given the high karyotype (Musilova, et al. 2013) and sequence (Jónsson, et al. 2014) identity with the Burchell’s zebra, we mapped the reads on the horse genome (EquCab3.0) and on the Equus_quagga_HiC assembly. As for Burchell’s zebra, we obtained CENP-A enrichment peaks which correspond to satellite-free centromeres (Figure 3, Supplementary Table S1). These neocentromeres were identified on 13 out of the 23 chromosomes. Eleven of these chromosomes (1, 3, 4, 5, 6, 9, 10, 11, 15, 16, and X) are meta- or sub-metacentric while two of them (19 and 20) are acrocentric. Eight satellite-free centromeres (5, 9, 10, 11, 15, 19, 20 and X) were identified on orthologous positions in both reference genomes (Figure 3 A and B). The centromeric domains of chromosomes 1, 4 and 16 were detected only in the horse reference genome suggesting that these loci are not properly assembled in the Equus_quagga_HiC scaffolds while the CENP-A binding domains of chromosomes 3 and 6 were detected only in the zebra assembly suggesting that the orthologous horse loci are highly rearranged.

**Figure 3.**
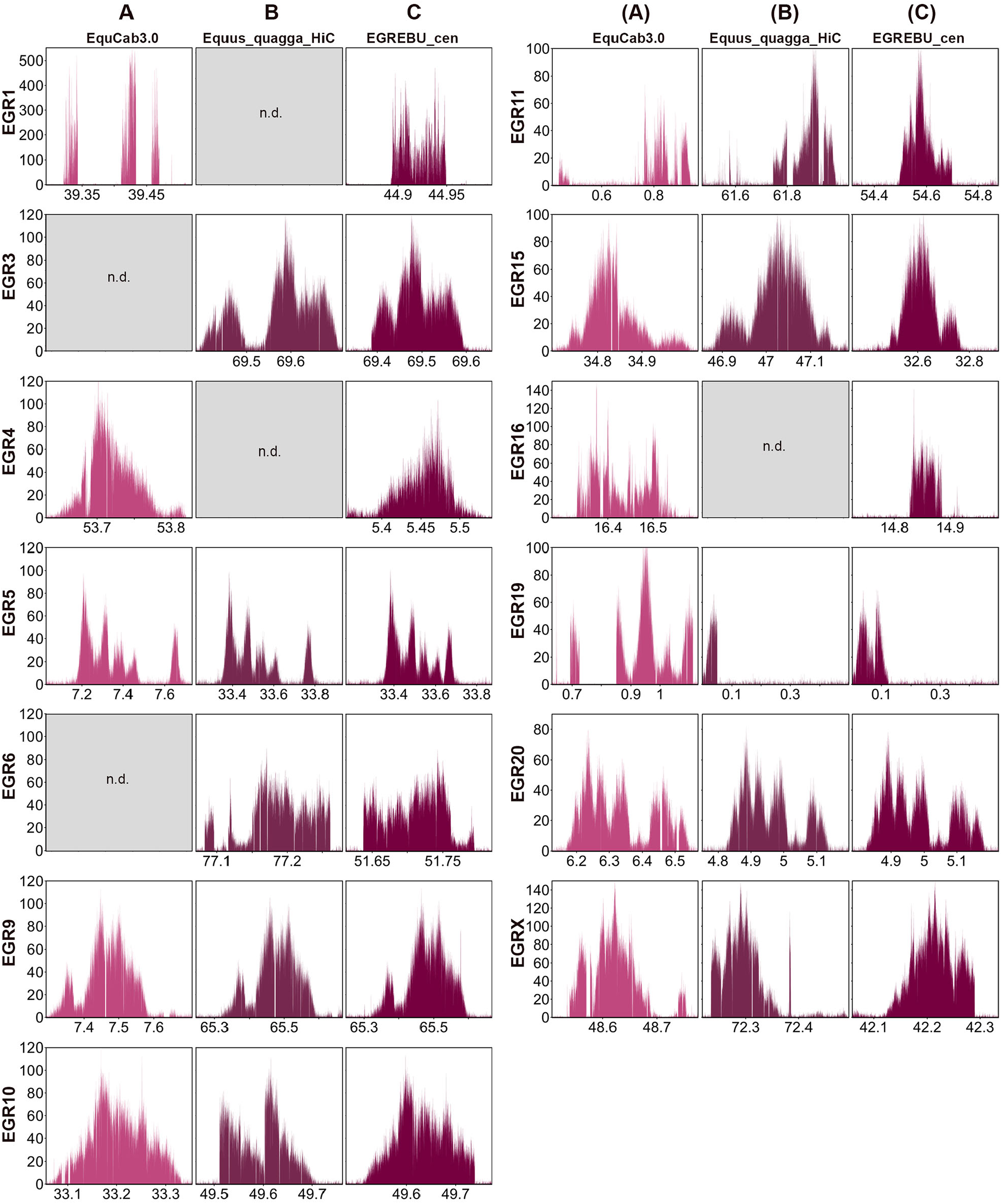
Satellite-free centromeres in Grevy’s zebra. ChIP-seq reads from primary fibroblasts of Grevy’s zebra were mapped on EquCab3.0 (A), Equus_quagga_HiC (B) and EGR_EBU_cen (C). CENP-A enriched domains are visualized as peaks. The y-axis reports the normalized read count whereas the x-axis reports the coordinates on the reference genome. For each chromosome, genomic windows of the same length are plotted.

Several enrichment peaks display irregular shapes due to sequence variation between the Grevy’s zebra and the reference genomes. To obtain a more precise reference for the centromeric loci of the Grevy’s zebra, the sequence corresponding to the 13 CENP-A binding peaks was assembled from our Illumina reads (see Materials and Methods) and contigs spanning between 60 and 370 kb were obtained (accession numbers: OM643415-OM643427). Using the approach previously applied by us to the donkey genome (Nergadze, et al. 2018), we constructed a chimeric reference genome by replacing the Equus_quagga_cen contigs with the othologous newly assembled Grevy’s zebra sequences (Supplementary Table S4). The result was a virtual hybrid reference genome, called EBU_EGR_cen, that was then used for mapping ChIP-seq reads to obtain refined CENP-A enrichment peaks (Figure 3C, Supplementary Table S5). As shown in Figure 3 C, several peak profiles were improved compared to those obtained using EquCab3.0 or Equus_quagga_HiC as reference genomes.

### Satellite-free centromeres with tandem repetitions

Similar to what previously described for some donkey satellite-free centromeres (Nergadze, et al. 2018), two Burchell’s zebra (EBU17 and EBU18) and two Grevy’s zebra (EGR1 and EGR16) centromeres display enrichment peaks with a spike-like shape (Figures 1C and 3C). We previously proved that, in the donkey, such peaks correspond to centromeres characterized by tandem repetitions of a sequence that was single-copy in the horse reference genome (Nergadze, et al. 2018). In particular, the EBU17 centromere has the same shape and resides in the same position of the centromere of donkey chromosome 16, which was previously shown to contain tandem repetitions (Nergadze, et al. 2018). Thus, we hypothesized that, also in the two zebras, the shape of such peaks indicates the presence of centromeric loci with DNA amplification. These loci could not be entirely assembled because of their repetitive nature.

To confirm the presence of tandem repetitions at these centromeres we analyzed the Burchell’s zebra input reads mapped on the Equus_quagga_cen reference genome and the Grevy’s zebra input reads mapped on the EBU_EGR_cen reference genome. The peaks shown in Figure 4A and C demonstrate that, at the genomic loci corresponding to the satellite-free centromeres of EBU17, EBU18, EGR1 and EGR16, multiple copies of the underlying genomic sequences are present. To approximately quantify the copy number of these repeats we compared the number of input reads mapping at the two Burchell’s zebra loci with the average genome coverage. As shown in Figure 4B, the number of reads at EBU17 and EBU18 centromeres is about 12.5 and 9.5 times, respectively, compared to genome average. The same analysis was carried out for the Grevy’s zebra loci and the number of reads at the EGR1 and at the EGR16 centromeres is about 8.3 and 2.8 times, respectively, compared to genome average.

**Figure 4.**
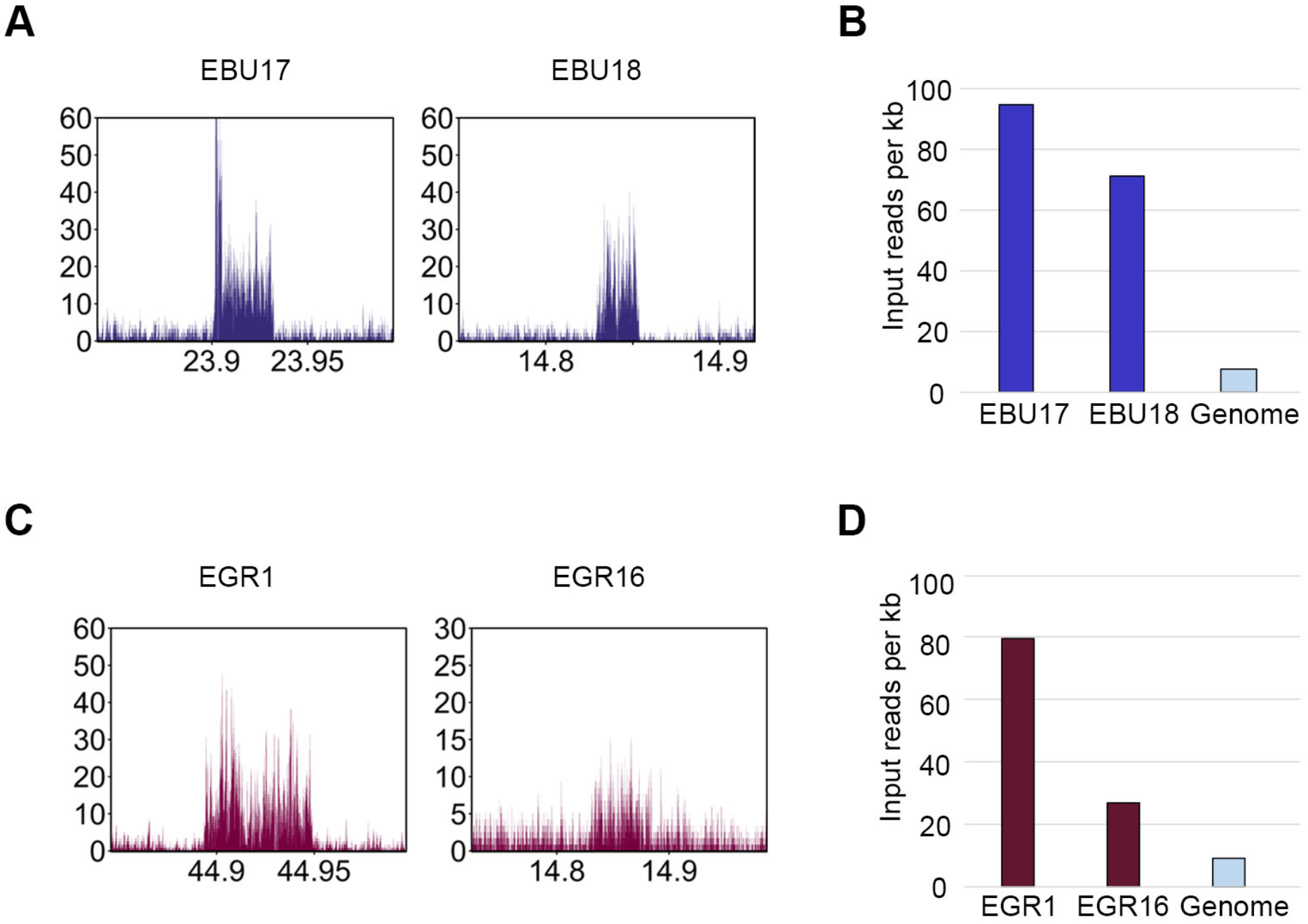
DNA sequence amplification at Burchell’s zebra and Grevy’s zebra satellite-free centromeres. A) Profiles of input (not immunoprecipitated) reads from Burchell’s zebra at EBU17 and EBU18 centromeric regions. The y-axis reports the RPKM count whereas the x-axis reports the coordinates on the reference genome. B) Input read counts per kilobase (y-axis) are shown for EBU17 and EBU18 neocentromeres and for the whole genome (Equus_quagga_cen). C) Profile of input reads from Grevy’s zebra at EGR1 and EGR16 centromeres. The y-axis reports the RPKM count whereas the x-axis reports the coordinates on the reference genome. D) Input read counts per kilobase (y-axis) are shown for EGR1 and EGR16 neocentromeres and the whole genome (EGR_EBU_cen).

The presence of sequence amplifications was also observed in the contigs that we assembled from our Illumina reads. In the EBU18, EGR1 and EGR16 CENP-A binding domains we detected the presence of repeated units of about 1.4, 5 and 3.1 kb, respectively (accession numbers: OM643411, OM643415 and OM643424). It is important to notice that EBU18 and EGR16 chromosomes are orthologous and their CENP-A binding domains are partially overlapping (Supplementary Table S1), suggesting that centromere formation and sequence amplification occurred before the separation of the two lineages.

### Sequence analysis of the satellite-free centromeres

DNA sequence features of the satellite-free centromeres of the two zebra species were compared with the average genome-wide values obtained from the Burchell’s zebra genome. The four centromeres containing tandem repetitions were excluded from this analysis because we cannot precisely define their complete underlying sequences. As shown in Supplementary Figure S2, in both species, the neocentromeres are significantly enriched in LINE-1 while depleted in LINE-2 elements, SINEs and DNA elements. On the other hand, the abundance of LTRs does not differ from the genome average. Given the enrichment in LINE-1 retrotransposons, we examined the sequence of these elements to test whether they are still active. The LINE-1 sequences identified at CENP-A binding domains seem transcriptionally silent being devoid of 3’ poly-A tails, often truncated and poorly conserved (data not shown) (Doucet, et al. 2015; Ivancevic, et al. 2016).

Finally, Burchell’s and Grevy’s zebra centromeres showed 37% and 36.6% GC content, respectively, which are lower than the genome-wide average of 41.43%. These differences are statistically significant proving that these centromeres are AT-rich (Supplementary Figure S2).

### Mechanisms of satellite-free centromeres formation in Burchell’s zebra

With the goal of determining the mechanisms of neocentromere formation during evolution, we carried out a comparative analysis between zebra chromosomes harboring satellite-free centromeres and the orthologous elements from the horse, here used as outgroup. Indeed, the horse is considered the closest extant species to the equid ancestor and all the centromeres, with the only exception of the one on chromosome 11, are satellite-based. Eight of the chromosomes (4, 6, 7, 8, 9, 10, 12 and 15) containing a satellite-free centromere were originated by fusion of ancestral chromosomes (Figure 5A). In all these chromosomes the ancestral centromeres were inactivated. To clarify the relationship between fusion and neocentromere formation, we measured the distance between each satellite-free centromere and the fusion point. To this purpose, since in the horse, as in the equid ancestor, the elements involved in the fusion are separated, we aligned the chromosomes from the Equus_quagga_cen assembly with the orthologous chromosomes from the EquCab3.0 horse assembly. Following this procedure, we were able to determine the coordinates of the fusion regions (Supplementary Table S6). These regions correspond to sequences placed on specific chromosomes of the Equus_quagga_cen assembly, range in size between about 5 kb and 1.5 Mb, and are not present as such in the horse genome assembly. A detailed description of all fusion events, as depicted in Figure 5A, and of the fusion regions is reported below.

**Figure 5.**
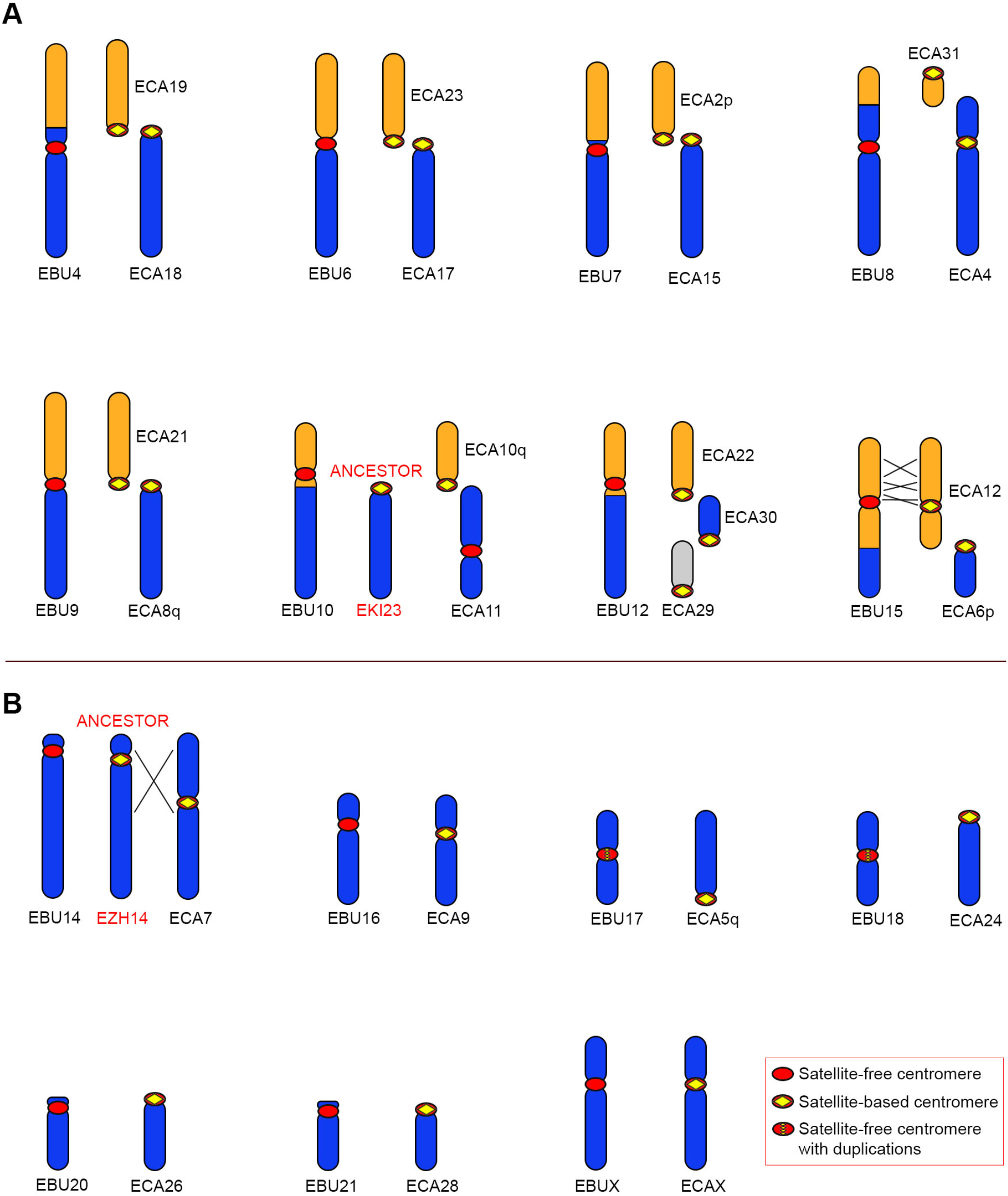
Comparison between Burchell’s zebra chromosomes with satellite-free centromeres and the orthologous horse chromosomes. For each Burchell’s zebra chromosome carrying a satellite-free centromere the corresponding orthologous horse chromosomes are shown. For EBU10 and EBU14, the ancestral chromosomes, represented by kiang chromosome 23 (EKI23) and Hartmann’s zebra chromosome 14 (EZH14), respectively, are shown. Inverted segments are indicated with crossed lines. A) EBU chromosomes deriving from fusions. B) EBU chromosomes corresponding to entire horse chromosomes or chromosome arms.

EBU4 is a metacentric chromosome deriving from the Robertsonian fusion of two ancestral acrocentric elements that correspond to horse chromosomes 18 and 19 (Figure 5A). The satellite-free centromeric region of EBU4 is orthologous to a non-centromeric locus on ECA18 and is located at about 6.5 Mb from the fusion region (Supplementary Table S6) suggesting that a centromere repositioning event moved the zebra centromere far from the fusion point. From these data it is not possible to determine whether centromere repositioning occurred before, during or after the fusion event. Interestingly, on the fusion region, we detected the presence of an array of 2PI, a satellite that we previously localized at most horse pericentromeres (Piras, et al. 2010; Cerutti, et al. 2016), of transposable elements and duplicons shared by different horse pericentromeres. All these sequences likely represent relics of the sequences associated to the ancestral centromeres which were involved in the Robertsonian fusion and are now inactive.

EBU6 derived from a fusion between ancestral acrocentrics orthologous to horse chromosomes 23 and 17 (Figure 5A). The CENP-A binding domain of EBU6 lays in a region that is unique in the Burchell’s zebra assembly but shares up to 80% identity with different horse pericentromeric sequences. This is the reason why this CENP-A binding domain was only detected when the ChIP-seq reads were mapped on the Burchell’s zebra genome while it was not detected when EquCab3.0 was used as reference (Figure 1). The organization of this fusion region is depicted in Figure 6A and shows that the CENP-A binding peak lays on a sequence sharing high identity with several horse pericentromeres, including the one from ECA 23 (light orange line in Figure 6A). The grey region, which contains the fusion point, is a highly rearranged 52 kb sequence with no good alignment with ECA23 nor ECA17 pericentromeres. On the other side of the grey region, a sequence sharing high identity with the pericentromere of ECA17 was detected (light blue line in Figure 6A). In conclusion, in EBU6, the CENP-A binding domain resides in the fusion region, within sequences deriving from an ancestral pericentromere. In Figure 6A the position of the CENP-A binding peak on Grevy’s zebra chromosome 6 is also reported and will be described in the next paragraph.

**Figure 6.**
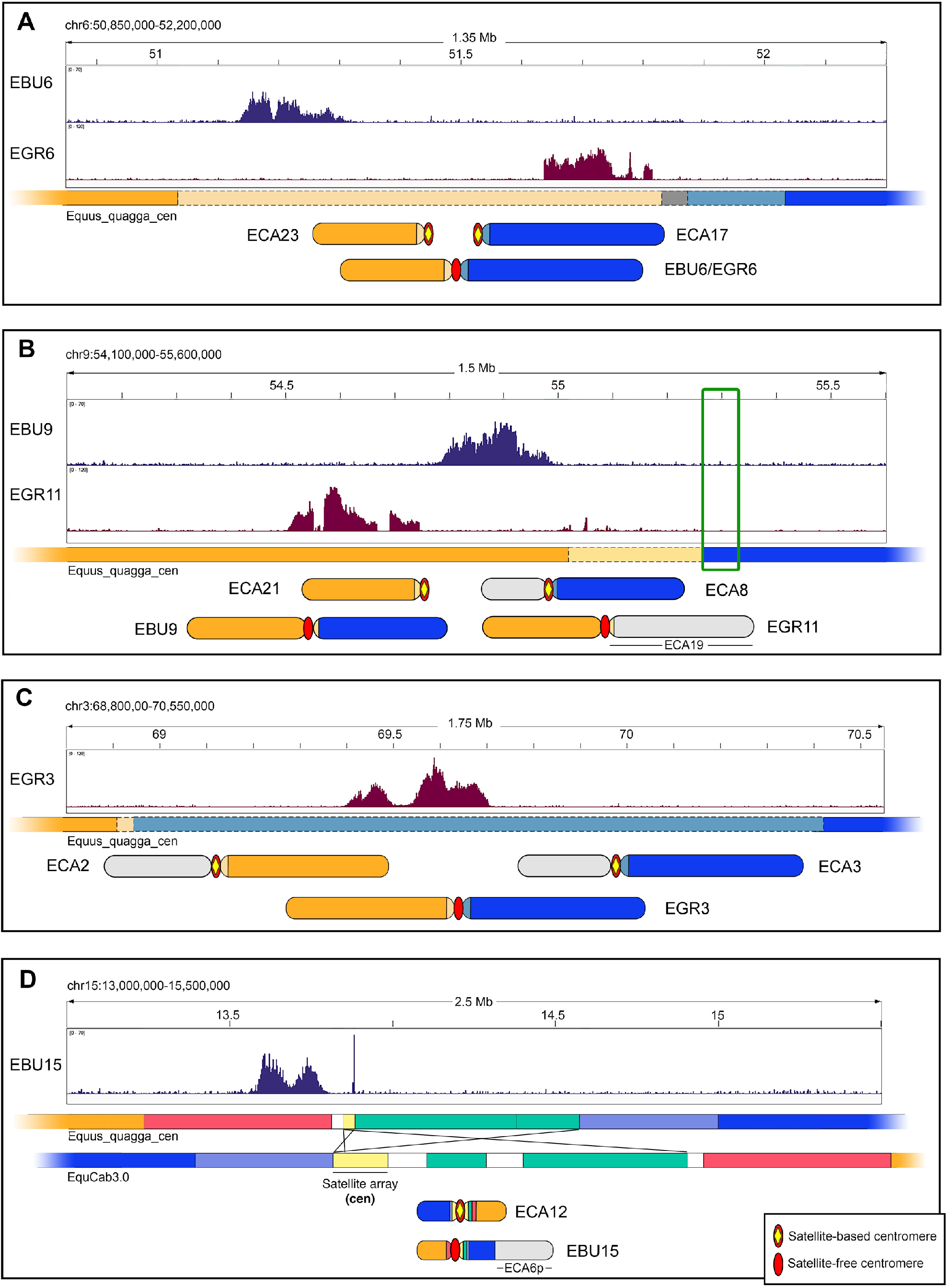
Sequence organization of satellite-free centromeric regions deriving from chromosomal rearrangements. A) The CENP-A binding peaks of EBU6 and EGR6 on the Equus_quagga_cen reference are shown on top. The colored bar schematically represents this Equus_quagga_cen genomic region with colors referring to orthologous sequences in the EquCab3.0 horse genome: orange, orthology with ECA23; blue, orthology with ECA17; light orange and light blue, orthology with horse pericentromeres, including ECA23 and ECA17, respectively; grey, orthology with several horse pericentromeric sequences not including ECA23 and ECA17. The zebra submetacentric chromosomes deriving from the fusion between ancestral elements corresponding to ECA23 (orange) and ECA17 (blue) are sketched on the bottom. B) The CENP-A binding peaks of EBU9 and EGR11 on the Equus_quagga_cen reference are shown on top. The colored bar schematically represents this Equus_quagga_cen genomic region with colors referring to orthologous sequences in the EquCab3.0 horse genome: orange, orthology with ECA21; blue, orthology with ECA8q; light orange, orthology with horse pericentromeres, including ECA21; the green box indicates satellite DNA. The EBU9 submetacentric chromosome, deriving from the fusion between ancestral elements corresponding to ECA21 (orange) and ECA8q (blue), together with the EGR11 submetacentric chromosome, deriving from the fusion between ancestral elements corresponding to ECA21 (orange) and ECA19 (light grey), are sketched on the bottom. The ECA8 p element, which did not participate in this fusion, is also in light grey. C) The CENP-A binding peak of EGR3 on the Equus_quagga_cen reference is shown on top. The colored bar schematically represents this genomic region with colors referring to orthologous sequences in the EquCab3.0 horse genome: orange, orthology with ECA2q; blue, orthology with ECA3q; light orange and light blue, orthology with horse pericentromeres, including ECA2q and ECA3q, respectively. The zebra submetacentric chromosome deriving from the fusion between ancestral elements corresponding to ECA2q (orange) and ECA3q (blue) are sketched on the bottom. ECA2p and ECA3p, which did not participate in this fusion, are in light grey. D) The CENP-A binding peak of EBU15 on the Equus_quagga_cen reference is shown on top. The colored bars represent the inverted regions in Equus_quagga_cen and in EquCab3.0. Colors mark orthologous segments between Burchell’s zebra and horse genomes. Crossed lines indicate inverted segments. Satellite arrays from the ECA12 pericentromere, which are partially assembled in EquCab3.0 reference, are in light yellow.

EBU7 derived from the fusion of the ancestral acrocentrics corresponding to ECA2p and ECA15 (Figure 5A). Its CENP-A binding domain lays in a region orthologous to a non-centromeric locus of ECA15 located 2 Mb away from the fusion region, where the relics of ancestral pericentromeric sequences, including blocks of 2PI and 137sat (Piras, et al. 2010; Nergadze, et al. 2014; Cerutti, et al. 2016) could be observed (Supplementary Table S6). Given the distance between the CENP-A binding domain and the fusion point, we hypothesize that the neocentromere of EBU7, similarly to EBU4, could be the result of a centromere repositioning event which occurred before, during or after the fusion.

EBU8 derived from a telomere-telomere fusion between elements orthologous to the acrocentric ECA31 and the submetacentric ECA4 (Figure 5A). At cytogenetic level, the centromere of EBU8 appears at the same position of the ECA4 centromere, however, at the molecular level, the CENP-A binding domain of EBU8 corresponds, in the horse, to a non-centromeric locus which is located at a distance of about 500 kb from 2PI pericentromeric satellites on ECA4. These observations suggest that the EBU8 centromere is repositioned relative to the ancestral ECA4 derived element. At the fusion site, located at 24 Mb from the CENP-A binding domain, no pericentromeric-type sequences were detected (Supplementary Table S6) and, although EBU8 derived from a telomere-telomere fusion, no telomeric repeats were detected either.

EBU9 resulted from the centric fusion between the ancestral segments corresponding to ECA21 and ECA8q (Figure 5A). The organization of this fusion region is depicted in Figure 6B. The EBU9 CENP-A binding domain is orthologous to a satellite-free non-centromeric locus of ECA21 (orange line in Figure 6B) and immediately adjacent to a region sharing high identity with ECA21 pericentromeric sequences (light orange line in Figure 6B). The fusion point is located at about 260 kb from the CENP-A binding domain, at the junction between the pericentromeric sequences derived from the ECA21 (light orange) and the ECA8q (blue in Figure 6B) ancestral elements. Moreover, in the junction region, 2PI and 137sat satellites (boxed in Figure 6B) can still be detected as traces of the ancient pericentromeres (Supplementary Table S6). Similar to EBU6, this satellite-free centromere resides very close to the fusion point. In Figure 6B the position of the CENP-A binding peak on Grevy’s zebra chromosome 11 is also reported and will be described in the next paragraph.

EBU10 derived from the fusion between ancestral elements corresponding to horse chromosomes 10q and 11 (Figure 5A). We previously demonstrated that horse chromosome 11 is totally deprived of satellite repeats and its centromere emerged from repositioning (Wade, et al. 2009; Piras, et al. 2010). Therefore, as shown in Figure 5A, ECA11 does not represent the ancestral configuration which is instead retained by the acrocentric chromosome 23 from *E. kiang* (Musilova, et al. 2013). EBU10 derived from a centric fusion and by a repositioning event that brought the centromere on the region orthologous to ECA10q. At the fusion site, that is located 4 Mb away from the CENP-A binding domain, arrays of 2PI and 137sat satellites were found (Supplementary Table S6).

EBU12 resulted from the fusion of three acrocentric chromosomes corresponding to horse chromosomes 22, 30 and 29 (Figure 5A). Its CENP-A binding domain corresponds to a non-centromeric locus on ECA22 and is located 10 Mb away from the ECA22/ECA30 fusion site, suggesting that this neocentromere emerged from repositioning. Also in this case, the fusion region contains arrays of the 2PI pericentromeric satellite (Supplementary Table S6).

EBU15 derives from the fusion between ancestral elements corresponding to horse chromosomes 12 and 6p (Figure 5A). The organization of this fusion region is depicted in Figure 6D. The CENP-A binding domain of EBU15 is orthologous to a non-centromeric horse locus which is located at a distance of about 1.1 Mb from the pericentromeric satellite of ECA12 (Figure 6D). The p-arm of EBU15 and the q-arm of ECA12 are not colinear but several rearrangements, including two previously undescribed inversions, occurred (Figure 5A). In addition, in the region immediately surrounding the centromere, two relatively small inversions differentiate the Burchell’s zebra and the horse genomes (Figure 6D). Interestingly, one of the breakpoints of this complex rearrangement falls exactly at the border of the satellite-free CENP-A binding domain, and contains pericentromeric 2PI repeats (yellow in Figure 6D). It is tempting to speculate that this break was involved in the inactivation of the ancestral centromere and in the formation of the new one.

In Figure 5B, the Burchell’s zebra chromosomes which carry satellite-free centromeres and did not derive from fusions (chromosomes 14, 16, 17, 18, 20, 21 and X) are depicted. They correspond to entire horse chromosomes or chromosome arms.

In the case of EBU14, previous cytogenetic comparative maps showed that the ancestral configuration is not represented by the horse chromosome 7 but rather by the Hartmann’s zebra chromosome 14 (Musilova, et al. 2013). Indeed, EBU14 is colinear with the ancestral chromosome but carries a repositioned satellite-free centromere while ECA7 derived from a pericentromeric inversion.

EBU16 and EBUX are submetacentric chromosomes entirely colinear with ECA9 and ECAX, respectively. In a previous cytogenetic study, the two Burchell’s zebra centromeres were considered at the same position of the horse ones (Musilova, et al. 2013). Here, sequence analysis allowed us to demonstrate that EBU16 and EBUX carry satellite-free centromeres which are orthologous to horse non centromeric loci (Supplementary Table S1).

EBU17 and EBU18 are entirely colinear with ECA5q and with the acrocentric chromosome ECA24, respectively (Piras, et al. 2009; Musilova, et al. 2013). Their centromeres moved from the ancestral terminal position to a new position void of satellite DNA, leading to the formation of metacentric chromosomes. Finally, from previous cytogenetic analysis (Musilova, et al. 2013) EBU20 and EBU21 were considered identical to their horse acrocentric orthologs, ECA26 and ECA28, respectively. Here we found that a satellite-free centromere is present on these two Burchell’s zebra chromosomes and that their CENP-A binding domains are located 4.9 and 1.7 Mb away from their p-arm terminus, respectively (Supplementary Table S3). The horse loci orthologous to these two neocentromeres do not contain functional centromeres. Satellite repeats, remnants of the ancestral satellite-based centromeres, are now located on the short arm (Piras, et al. 2010). Therefore, the short arms of EBU20 and EBU21 are a few Mb longer than the horse ones.

### Mechanisms of satellite-free centromeres formation in Grevy’s zebra

We investigated the mechanisms of neocentromere formation in Grevy’s zebra. As mentioned above, a reference genome was not available for this species but, given the high karyotype (Musilova, et al. 2013) and sequence (Jónsson, et al. 2014) identity with Burchell’s zebra, we used our Burchell’s zebra genome assembly (Equus_quagga_cen) and the hybrid genome (EBU_EGR_cen) as reference.

In Grevy’s zebra, chromosomes 1, 3, 4, 5, 6, 9, 10 and 11 were originated by fusions and carry a satellite-free centromere (Figure 7A). In all these chromosomes the ancestral centromeres were inactivated. At the cytogenetic level, chromosomes 1, 3, 5, 6 and 10 are colinear with Burchell’s zebra chromosomes 1, 3, 5, 6 and 10, respectively, suggesting that these fusion events occurred in the common ancestor of the two species (Musilova, et al. 2013). Differently, Grevy’s zebra chromosomes 4, 9 and 11 derived from lineage-specific fusion events. A description of all fusion events, as depicted in Figure 7A, is reported below.

**Figure 7.**
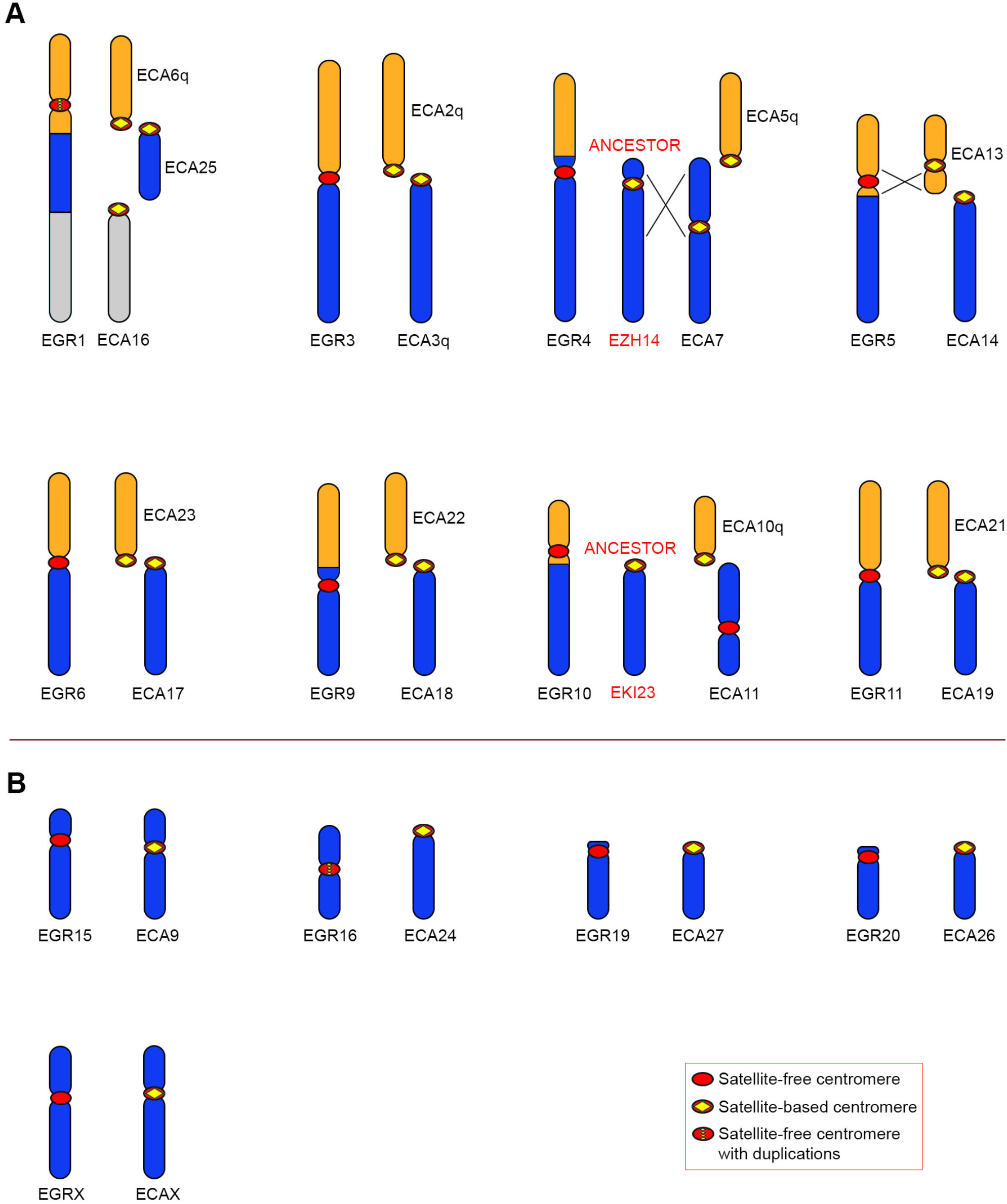
Comparison between Grevy’s zebra chromosomes with satellite-free centromeres and the orthologous horse chromosomes. For each Grevy’s zebra chromosome carrying a satellite-free centromere, the corresponding orthologous horse chromosomes are shown. For EGR4 and EGR10, the ancestral chromosomes, represented by Hartmann’s zebra chromosome 14 (EZH14) and kiang chromosome 23 (EKI23), respectively, are reported. Inverted segments are indicated with crossed lines. A) EGR chromosomes deriving from fusions. B) EGR chromosomes corresponding to entire horse chromosomes or chromosome arms.

EGR1 derived from the fusion of ancestral elements corresponding to horse chromosomes 6q, 25 and 16 (Figure 7A). Interestingly, this chromosome, which is characterized by a satellite-free centromere, is orthologous to EBU1 whose centromere, being satellite-based, could not be precisely mapped. At the cytogenetic level, the EBU1 and EGR1 centromeres appear at the same position but, through sequence analysis, we showed that the CENP-A binding domain of EGR1 is located at a distance of 10 Mb from the fusion region between ECA6q and ECA25 (Supplementary Table S6) suggesting that a repositioning event moved this centromere far from the fusion point. The sequence of this neocentromere, which is characterized by the presence of amplifications, could not be found in the Burchell’s zebra genome assembly (Figure 3). EGR3 derived from a centric fusion involving two ancestral acrocentric chromosomes orthologous to horse chromosomes 2q and 3q (Myka, et al. 2003) and is colinear with EBU3. The organization of this fusion region is depicted in Figure 6C. While EBU3 contains a satellite-based centromere, the CENP-A binding domain of EGR3 resides on a sequence that is unique in Burchell’s zebra but shares high identity with several horse pericentromeres, including the one from horse chromosome 3q (Figure 6C). This is the reason why this neocentromere was detected in the Burchell’s zebra assembly but not in the EquCab3.0 reference genome (Figure 3). The fusion point lays about 500 kb upstream of the centromeric domain where a good alignment with ECA2q pericentromeric sequences was observed (light orange in Figure 6C). The presence of the CENP-A binding domain within the ancestral pericentromeric sequences suggests that the centric fusion contributed to neocentromere formation.

EGR4 derived from the fusion of two ancestral elements corresponding to horse chromosome 5q and Hartmann’s zebra chromosome 14, which is orthologous to Burchell’s zebra chromosome 14 (Figure 5A and 7A). The neocentromere of EGR4 lays in the same position of the one on EBU14 (Supplementary Table S1), suggesting that a centromere repositioning event occurred in the common ancestor of the two zebra species before the fusion with the ECA5q element in the Grevy’s zebra lineage.

EGR5 is orthologous to EBU5 (Musilova, et al. 2013), however, although the centromere of EGR5 is satellite-free, the one from EBU5 is satellite-based (Piras, et al. 2010). EGR5 was originated by the fusion between ancestral elements corresponding to horse chromosomes 13 and 14. From previous cytogenetic analysis, an inversion was identified in the EGR5 segment orthologous to ECA13 (Musilova, et al. 2013). Our sequence analysis confirmed that the EGR5 neocentromere is contained in the inverted region (Figure 7A). The EGR5 CENP-A binding domain is located 4.8 Mb away from the fusion site which contains 2PI satellite arrays (Supplementary Table S6).

EGR6, deriving from the centric fusion between elements corresponding to ECA17 and ECA23 (Figure 7A), is orthologous to EBU6 which was described above (Figure 5A). Notably, the CENP-A binding domain of EGR6 is only 13 kb away from the 52 kb region containing the fusion point. As shown in Figure 6A, the CENP-A binding domains of the neocentromeres on EGR6 and EBU6 are located only 278 kb apart on the reference genome (Figure 6A and Supplementary Table S1).

EGR9 is partially orthologous to EBU4 (Musilova, et al. 2013) (Figure 5A and 7A). These two chromosomes derive from different fusion events both involving an ancestral element corresponding to horse chromosome 18. In EGR 9 this element was fused with an element orthologous to horse chromosome 22 (Figure 7A) while in EBU4 it was fused with an element orthologous to horse chromosome 19 (Figure 5A). In EGR9 and in EBU4 the satellite-free centromere is localized in the same genomic region on the ECA18 derived element (Supplementary Table S1) at about 6.5 Mb from the fusion region.

EGR10 is orthologous to EBU10 (Musilova, et al. 2013) (Figure 5A and 7A). These two chromosomes carry their satellite-free centromeres in the same genomic locus at about 4 Mb from the fusion site (Supplementary Table S1 and S6), suggesting that chromosome fusion and neocentromere formation occurred in their common ancestor.

EGR11 is partially orthologous to EBU9 (Musilova, et al. 2013) (Figure 5A and 7A). EGR11 and EBU9 derive from different fusion events involving an ancestral element corresponding to ECA21. The organization of this fusion region is depicted in Figure 6B. In EGR11 this element was fused with an element corresponding to ECA19 (Figure 7A) while in EBU9 it was fused with an element corresponding to ECA8q (Figure 5A). The EGR11 CENP-A binding domain is orthologous to a satellite-free non-centromeric locus on ECA21 and lays in the same chromosomal locus of the satellite-free centromere of EBU9 (Figure 6B). These neocentromeres are adjacent to the fusion region (Figure 6B).

The remaining Grevy’s zebra chromosomes carrying a neocentromere (EGR15, 16, 19, 20 and X) correspond to entire horse chromosome or chromosome arms and are entirely colinear with them (Figure 7B). All these centromeres reside in genomic regions orthologous to non-centromeric regions in the horse. The CENP-A binding domains of EGR15, 16, 20 and X are located in the same genomic position of the ones of their Burchell’s zebra orthologs (EBU16, 18, 20 and X), suggesting that they were originated before the separation of the two lineages.

## DISCUSSION

### Identification of neocentromeres in Burchell’s and Grevy’s zebra

The DNA Zoo team recently released the first genome assembly for the Burchell’s zebra (Equus_quagga_HiC). This genome sequence contains scaffolds that correspond to entire zebra chromosomes but lack chromosome assignment. In this work, thanks to the availability of sub-chromosomal comparative maps between the Burchell’s zebra and the horse (Musilova, et al. 2013), we improved the DNA Zoo assembly by assigning chromosome numbers to scaffolds and adjusting the direction of some chromosomes. To further improve the Burchell’s zebra reference genome we replaced the regions surrounding the 15 satellite-free centromeres with sequences that we *de novo* assembled from our ChIP-seq reads.

The new assembly was also used as reference to identify the 13 satellite-free centromeres of Grevy’s zebra, a species closely related to Burchell’s zebra but still lacking a reference genome.

This work allowed us to demonstrate, at the DNA sequence level, that Burchell’s zebra and Grevy’s zebra are characterized by an extraordinary high number of centromeres completely devoid of satellite DNA. Indeed, as for the donkey (Nergadze, et al. 2018), more than half of their chromosomes have satellite-free centromeres, further proving that the absence of satellite DNA at centromeric domains is compatible with genome stability and species survival.

Using this approach, we could not obtain ChIP-seq peaks from the satellite-based centromeres because their corresponding reads could not be mapped on specific chromosomes.

Previous work from our laboratory demonstrated that the position of satellite-free centromeric domains can vary in the horse and donkey populations (Purgato, et al. 2015; Nergadze, et al. 2018; Cappelletti, et al. 2019). Here, although we analyzed only one individual from each species, we were able to observe, in a few cases, two well separated CENP-A binding domains, corresponding to different epialleles on the two homologs. This observation confirms that the phenomenon of centromere sliding that we previously observed in horse and donkey is occurring also in the two zebras.

Seven Burchell’s zebra centromeres (from EBU6, EBU7, EBU8, EBU9, EBU16, EBU17, EBU18 chromosomes) and nine Grevy’s zebra centromeres (from EGR1, EGR3, EGR4, EGR5, EGR6, EGR10, EGR11, EGR15, EGR16 chromosomes) were shown to lack satellite DNA at the FISH resolution level (Piras, et al. 2010). Here we confirmed, at the DNA sequence level, that they are completely satellite-free. However, not all the satellite-free centromeres identified in the present work coincide with those described in our previous cytogenetic analysis (Piras, et al. 2010). The first discrepancy concerns several chromosomes whose primary constrictions seemed to be associated to satellite repeats (EBU4, EBU10, EBU12, EBU14, EBU15, EBU20, EBU21, EBUX, EGR9, EGR19, EGR20) (Piras, et al. 2010). In the present work, we found that the CENP-A binding domains of these chromosomes lay on single-copy regions. We propose that these satellite-free neocentromeres formed relatively close to highly repetitive tandem repeats cytogenetically coinciding with primary constrictions. The satellite repeats previously observed by FISH may correspond to ancestral centromeric domains that are now inactive while the CENP-A binding domains lay on single-copy regions. These observations indicate that the centromeric function has shifted away from a satellite repeat containing locus to a satellite-free new position and confirm that satellite DNA sequences are not sufficient nor necessary for specifying centromere function.

In addition, at seven Grevy’s zebra chromosomes (EGR2, EGR8, EGR13 EGR14, EGR17, EGR18, EGR22), although no satellite FISH signals were detected in our previous cytogenetic analysis, we did not identify, in the present study, satellite-free centromeres by ChIP-seq. It is possible that, at these “elusive” centromeres, short or chromosome specific tandem arrays, undetectable at the FISH resolution level, are present. Alternatively, since a Grevy’s zebra reference genome is not available, it is possible that their CENP-A binding domains lay on single copy regions that may be absent both in the Burchell’s zebra and in the horse assemblies suggesting that the number of Grevy’s zebra neocentromeres may be underestimated.

### DNA sequence composition of neocentromeres

The extraordinary high number of sequenced satellite-free centromeres allowed us to investigate whether any conserved sequence feature is present at these genomic regions. As previously shown for the donkey satellite-free centromeres (Nergadze, et al. 2018), the Burchell’s and Grevy’s zebra centromeres are LINE-1 rich. Although not universal, the presence of retroelements is common at the centromeres of many mammals, insects, plants and fungi (Plohl, et al. 2008; Longo, et al. 2009; Klein and O’Neill 2018; Yadav, et al. 2018; Chang, et al. 2019) while it is still under debate whether these elements might promote genome instability of these regions. Although, at some sporadic human neocentromeres, LINE elements can be active (Chueh, et al. 2005; Marshall, et al. 2008; Chueh, et al. 2009; Barra and Fachinetti 2018), in the evolutionary satellite-free centromeres described here, the LINE-1 elements seem to be inactive.

It is well described that LINE-1 elements are preferentially inserted into AT-rich sequences which display low nucleosome occupancy (Sultana, et al. 2019). AT richness is a typical feature of centromeres in a number of organisms (Clarke and Carbon 1985; Marshall, et al. 2008; Talbert and Henikoff 2020) and might favor the adoption of a non-B DNA configuration which is usually found at centromeres (Kasinathan and Henikoff 2018; Talbert and Henikoff 2020). It was recently shown that AT-rich exogenous DNA is also capable of functioning as a centromere in the model organism *Schizosaccharomyces pombe* (Barbosa, et al. 2022). As previously shown for the 16 satellite-free donkey centromeres (Nergadze, et al. 2018), the neocentromeres of the two zebras are AT-rich as well suggesting that such sequence feature may favor CENP-A binding and other epigenetic modifications.

Sequence analysis revealed that, in two Burchell’s zebra (from EBU17 and EBU18) and in two Grevy’s zebra (from EGR1 and EGR16) neocentromeres, amplification of DNA sequences occurred. We previously proposed that evolutionary new centromeres, initially devoid of satellite DNA, can undergo a process of “maturation” during their evolution through the acquisition of satellite DNA (Piras, et al. 2010). In this view, the presence of duplications, that we previously observed also at a subset of donkey neocentromeres, has been interpreted as the first step of centromere “satellitization” (Nergadze, et al. 2018). At EBU18 and EGR16, which are orthologous and completely colinear, the CENP-A binding domains are partially overlapping on the reference genomes, suggesting that centromere formation and sequence amplification occurred before the separation of the two lineages. Although in the commonly accepted phylogenetic tree of equids (Jónsson, et al. 2014) the lineages of donkey and Burchell’s zebra are separated, EBU17 is orthologous and completely colinear with donkey chromosome 16 and their centromeres are both characterized by the presence of amplifications (Nergadze, et al. 2018). A possible explanation of this finding is that centromere repositioning occurred in a common ancestor of asses and zebras and that lineage sorting or chromosome rearrangement was then responsible for maintaining this centromere only in these two lineages. Alternatively, we can hypothesize that independent repositioning events occurred in the two lineages at a hotspot for neocentromere formation similar to what already observed for human clinical neocentromeres (Marshall, et al. 2008). Another observation supporting this hypothesis is the formation of clinical neocentromeres at positions corresponding to ancestral non-human primate centromeres, where presumably a latent centromere forming potentiality persisted (Ventura, et al. 2004; Capozzi, et al. 2009; Rocchi, et al. 2012).

### Conservation of centromeric domains

The Burchell’s and the Grevy’s zebras share 13 orthologous chromosomes which are identical at the cytogenetic level (Trifonov, et al. 2012; Musilova, et al. 2013). In the present work we demonstrated that in only one of the couples of orthologs (EBU2/EGR2) both centromeres are satellite-based while six out of these couples (EBU6/EGR6, EBU10/EGR10, EBU16/EGR15, EBU18/EGR16, EBU20/EGR20 and EBUX/EGRX) harbor a satellite-free centromere in the same locus, suggesting that neocentromere formation occurred in the common ancestor of the two species. These evolutionary new centromeres did not have the time to acquire the typical complexity of mammalian centromeres. This situation is consistent with the evolutive youth of these species, which emerged about 1.4 million years ago (Jónsson, et al. 2014). On the other hand, six orthologous pairs (EBU1/EGR1, EBU3/EGR3, EBU7/EGR7, EBU8/EGR8, EBU15/EGR14, EBU19/EGR19) are cytogenetically colinear but we could detect a satellite-free centromere in one of the two zebras only. The EBU19 and EGR19 orthologs are small chromosomes that were described as acrocentric (Musilova, et al. 2013). In EGR19, the centromere might have been repositioned very close to the terminal position while the ancestral satellite-based centromere was maintained in EBU19. All the other pairs of orthologs with divergent centromeres derived from fusion events. Two alternative explanations can be proposed to explain these observations: 1) Independent fusion events involving the same ancestral elements and inactivation of the old centromeres occurred in the two species. In this scenario, only one of the two fusion events was successively accompanied by the formation of a satellite-free centromere; 2) Chromosome fusion and neocentromere formation, occurred in the common ancestor of the two zebras but then centromere maturation took different routes in the two species generating new satellite repeats in one species only. In this scenario, for each pair of orthologous chromosomes, the centromere of a species acquired new satellite DNA sequences, reaching the typical complexity of mammalian centromeres, while in the other lineage the “immature” satellite-free centromere was maintained. The case of EBU1 and EGR1 may support the second possibility: in EBU1 a satellite-based centromere is present while in EGR1 a satellite-free centromere with amplified sequences may represent an intermediate maturation step.

There are other pairs of chromosomes, that are only partially colinear, in which both species carry a satellite-free neocentromere in the same genomic locus (EBU4/EGR9, EBU9/EGR11 and EBU14/EGR4). A possible explanation of this finding is that these neocentromeres appeared in a common ancestor and that lineage-specific fusion events occurred after neocentromere formation. Alternatively, two different Robertsonian fusions occurred in the two species and then satellite-free centromeres arose independently in the same “centromerization” hot spot.

### Mechanisms for satellite-free centromeres formation

The rapid equid evolution was characterized by extensive karyotype reshaping. Neocentromere formation and inactivation of old satellite-based centromere played a key role in this process. In horse and donkey, the mechanism for satellite-free centromere formation was the shift of the centromeric function from an ancestral terminal position to a new interstitial position not involving chromosome rearrangement (Carbone, et al. 2006; Wade, et al. 2009; Piras, et al. 2010; Nergadze, et al. 2018). In Burchell’s and Grevy’s zebras we observed several neocentromeres that were generated through this mechanism, however chromosome rearrangements, particularly chromosome fusions, were also involved.

Interestingly, the CENP-A binding domain often resides in the fusion regions, on pericentromeric sequences derived from the ancestral inactivated centromeres, strongly suggesting that the Robertsonian fusion directly triggered the formation of a neocentromere within the fusion region.

Our sequence analysis revealed that, in zebras, the centromeres of chromosomes derived from Robertsonian fusions are different from the ones previously described in mouse (Garagna, et al. 2001; Garagna, et al. 2014). Cytogenetic analysis showed that, in the mouse, the fusion region maintains long stretches of satellite repeats with an anti-parallel symmetry and an equal contribution of the two ancestral telocentric chromosomes in the formation of the new satellite-based centromere (Garagna, et al. 2001). On the contrary, the molecular analysis presented here showed that, in most zebra Robertsonian fusions, the new centromere is satellite-free while the ancestral satellite repeats are lost.

A peculiar neocentromere is the one on EBU15 where an inversion was probably involved in its formation. Indeed, this satellite-free centromere was seeded on an inversion breakpoint.

It has been suggested that centromere strength is directly correlated with the length of satellite arrays. According to this view, the fixation of fused chromosomes can result from meiotic drive and occur when the centromere of a bi-armed chromosome is “stronger” than the original centromeres, ending in its preferential segregation into the egg during female meiosis (Fishman and Saunders 2008; Chmátal, et al. 2014; Iwata-Otsubo, et al. 2017; Wei, et al. 2017; Kursel and Malik 2018; Talbert and Henikoff 2020). According to this model, extended satellite arrays would allow CENP-A nucleosome expansion resulting in the formation of a stronger kinetochore compared to short arrays. On the contrary, the results presented here show that, in zebras, neocentromeres completely devoid of satellite DNA are present in fusion chromosomes indicating that the contribution of repeated DNA in strengthening the centromere is not universal and that other genetic or epigenetic features may guarantee the stability of centromeres. It is also possible that factors such as small population size and bottle-neck during evolution may have contributed to the fixation of satellite-free centromeres in Robertsonian chromosomes (Trifonov, et al. 2008; Ransom and Kaczensky 2016).

## METHODS

### Cell lines

Primary fibroblast cell lines from Burchell’s zebra and Grevy’s zebra were previously described (Piras, et al. 2010). The cells were cultured in high-glucose DMEM medium supplemented with 20% fetal bovine serum, 2 mM L-glutamine, 1% penicillin/streptomycin and 2% non-essential amino acids. Cells were maintained at 37 °C in a humidified atmosphere of 5% CO2.

### ChIP-seq

Chromatin from primary fibroblasts was cross-linked with 1% formaldehyde, extracted, and sonicated to obtain DNA fragments ranging from 200 to 800 bp. Immunoprecipitation was performed as previously described (Nergadze, et al. 2018) by using an anti-CENP-A serum (Cappelletti, et al. 2019). Paired-end sequencing was performed through an Illumina HiSeq2500 platform by IGA Technology Services (Udine, Italy).

### Comparative genomic analysis

Pairwise alignments between whole genomes were performed with Chromeister (version 1.5a) (Pérez-Wohlfeil, et al. 2019) available at the European Galaxy web platform (https://usegalaxy.eu/) (Afgan, et al. 2016) using default parameters. The aligned genomes were the EquCab3.0 horse genome, the Equus_quagga_HiC Burchell’s zebra genome and the modified Burchell’s zebra genome we produced, Equus_quagga_cen. The Equus_quagga_HiC draft assembly, available at https://www.dnazoo.org/assemblies/Equus_quagga, was generated by the DNA Zoo consortium from short insert-size PCR-free DNA-Seq data using w2rap-contigger (Clavijo, et al. 2017), see (Dudchenko, et al. 2017; Dudchenko, et al. 2018) for details. The blood sample for *in situ* Hi-C preparation was donated by a female individual named Zena, and obtained from Nancy Nunke (Hearts & Hands Animal Rescue), Greg Barsh (Stanford, Hudson Alpha), Ren Larison (UCLA).

Pairwise alignments between single chromosomes were run locally using the same Chromeister version (https://github.com/estebanpw/chromeister) with default parameters. The resulting plots were used to evaluate chromosome orthologies and Burchell’s zebra chromosome orientations. Fine-scaled orthologies were resolved by BLAT-searching a sequence of interest from the Burchell’s zebra genome against the horse EquCab3.0 reference using UCSC Genome Browser (https://genome.ucsc.edu/index.html).

### Bioinformatic analysis of ChIP-seq data

Reads were aligned with paired-end mode to the reference genomes with Bowtie2 aligner (version 2.4.2) (Langmead, et al. 2009; Langmead and Salzberg 2012). Normalization of read coverage of the ChIP datasets against the input datasets was performed using bamCompare available in the deepTools suite (3.5.0 version) (Ramírez, et al. 2016) using RPKM (Reads Per Kilobase per Million mapped reads) normalization in subtractive mode. Normalization of read coverage of the input datasets for amplification analysis was performed using bamCoverage available in the deepTools suite (Ramírez, et al. 2016) using RPKM (Reads Per Kilobase per Million mapped reads) normalization. The resulting coverage files were visualized using Integrative Genomics Viewer (IGV) software (http://software.broadinstitute.org/software/igv/home). Peaks were obtained with the R software package Sushi (Phanstiel, et al. 2014). Peak calling was performed with MACS2 (version 2.2.7.1) (Zhang, et al. 2008; Feng, et al. 2012) using --broad option to identify large enriched genomic regions. We did not include in our analysis enriched regions containing satellite arrays. Enriched regions from Equus_quagga_HiC were BLAT-searched against EquCab3.0 using UCSC Genome Browser (https://genome.ucsc.edu/index.html) to confirm the correspondence with the orthologous enriched loci identified in the horse genome.

### Assembly of centromeric regions, improvement of the Burchell’s zebra reference genome and construction of a chimeric genome

The *de novo* assembly of centromeric regions from Burchell’s and Grevy’s zebras was performed using an iterative chromosome walking approach based on paired-end ChIP-seq reads, as previously described (Nergadze, et al. 2018).

To obtain a refined reference genome for Burchell’s zebra, we first run BLAT (v. 36) (Kent 2002) using the assembled contigs as query to identify their incomplete and/or misassembled counterparts in the Equus_quagga_HiC scaffolds. The incomplete and/or misassembled sequences were then removed and substituted with the newly-assembled centromeric contigs using SAMtools (version 1.11) (Danecek, et al. 2021). Finally, when the orientation of the Equus_quagga_HiC scaffolds identified through the Chromeister tool was reverse, we adjusted it using the Reverse.seq tools (Galaxy Version 1.39.5.0) (Schloss, et al. 2009) available at the Galaxy web platform (https://usegalaxy.org/) (Afgan, et al. 2016).

To obtain a more accurate reference for Grevy’s zebra, the regions orthologous to the newly-assembled contigs were identified by BLAT (v. 36) in the Equus_quagga_cen genome. These regions were removed and substituted with the Grevy’s zebra centromeric contigs to obtain a chimeric reference genome, as previously described (Nergadze, et al. 2018).

The resulting reference genomes were used to re-map ChIP-seq reads and obtain enrichment peaks, as described above.

### Sequence analysis of centromeric domains

The presence of DNA amplification at centromeres was evaluated by comparing the read counts per kilobase in the centromeric domains with the genome-wide average using input reads aligned to the corrected Burchell’s zebra reference genome or to the chimeric reference genome. Counts were obtained using SAMtools suite (version 1.11).

The analysis of the content in repetitive elements in centromeric domains was carried out with RepeatMasker (Galaxy Version 4.0.9) (http://www.repeatmasker.org) available at the Galaxy web platform. The same analysis was carried out on the entire Burchell’s zebra genome in order to obtain genome-wide values. Statistical significance was assessed by t-test using the VassarStats website by comparing the values of centromeric genomic regions with the values of genome scaffolds (http://vassarstats.net/). Centromeres containing DNA amplifications were excluded from this analysis.

## DATA AVAILABILITY

Raw sequencing data from this study have been submitted to the NCBI BioProject database (https://www.ncbi.nlm.nih.gov/bioproject/) under accession number PRJNA804871. De novo assembled centromeric regions of Burchell’s zebra and Grevy’s zebra from this study have been submitted to the NCBI Nucleotide database (https://www.ncbi.nlm.nih.gov/nucleotide/) under accession numbers OM643400-OM643427.

## ACKNOWLEDGEMENTS

We would like to thank Francesco Lescai for helpful suggestions on the bioinformatic analysis. This work was supported by the Italian Ministry of Education, University and Research (MIUR) [Dipartimenti di Eccellenza Program (2018-2022) - Dept. of Biology and Biotechnology “L. Spallanzani”, University of Pavia], PRIN Grant No. 2015RA7XZS_002 and from the Consiglio Nazionale delle Ricerche (CNR-Progetto Bandiera Epigenomica).

## AUTHOR CONTRIBUTIONS

E.G. conceived and supervised the study. E.C., F.M.P., S.G.N and E.G. contributed to experimental design. E.C., F.M.P. and S.G.N. carried out molecular and cell biology experiments and bioinformatic analyses. E.C., F.M.P., S.G.N. and E.G. wrote the manuscript. L.S., M.S., W.A.A., E.R. contributed to analyses. All authors read and approved the final manuscript.

## REFERENCES

Afgan E, Baker D, van den Beek M, Blankenberg D, Bouvier D, Čech M, Chilton J, Clements D, Coraor N, Eberhard C, et al. 2016. The Galaxy platform for accessible, reproducible and collaborative biomedical analyses: 2016 update. Nucleic Acids Res 44:W3–W10.

Allshire RC, Karpen GH. 2008. Epigenetic regulation of centromeric chromatin: old dogs, new tricks? Nat Rev Genet 9:923–937.

Barbosa AC, Xu Z, Karari K, Williams W, Hauf S, Brown WRA. 2022. Mutation and selection explain why many eukaryotic centromeric DNA sequences are often A + T rich. Nucleic Acids Res 50:579–596.

Barra V, Fachinetti D. 2018. The dark side of centromeres: types, causes and consequences of structural abnormalities implicating centromeric DNA. Nat Commun 9:4340.

Capozzi O, Purgato S, D’Addabbo P, Archidiacono N, Battaglia P, Baroncini A, Capucci A, Stanyon R, Della Valle G, Rocchi M. 2009. Evolutionary descent of a human chromosome 6 neocentromere: a jump back to 17 million years ago. Genome Res 19:778–784.

Cappelletti E, Piras FM, Badiale C, Bambi M, Santagostino M, Vara C, Masterson TA, Sullivan KF, Nergadze SG, Ruiz-Herrera A, et al. 2019. CENP-A binding domains and recombination patterns in horse spermatocytes. Sci Rep 9:15800.

Carbone L, Nergadze SG, Magnani E, Misceo D, Francesca Cardone M, Roberto R, Bertoni L, Attolini C, Francesca Piras M, de Jong P, et al. 2006. Evolutionary movement of centromeres in horse, donkey, and zebra. Genomics 87:777–782.

Cerutti F, Gamba R, Mazzagatti A, Piras FM, Cappelletti E, Belloni E, Nergadze SG, Raimondi E, Giulotto E. 2016. The major horse satellite DNA family is associated with centromere competence. Mol Cytogenet 9:35.

Chang CH, Chavan A, Palladino J, Wei X, Martins NMC, Santinello B, Chen CC, Erceg J, Beliveau BJ, Wu CT, et al. 2019. Islands of retroelements are major components of Drosophila centromeres. PLoS Biol 17:e3000241.

Chmátal L, Gabriel SI, Mitsainas GP, Martínez-Vargas J, Ventura J, Searle JB, Schultz RM, Lampson MA. 2014. Centromere strength provides the cell biological basis for meiotic drive and karyotype evolution in mice. Curr Biol 24:2295–2300.

Chueh AC, Northrop EL, Brettingham-Moore KH, Choo KH, Wong LH. 2009. LINE retrotransposon RNA is an essential structural and functional epigenetic component of a core neocentromeric chromatin. PLoS Genet 5:e1000354.

Chueh AC, Wong LH, Wong N, Choo KH. 2005. Variable and hierarchical size distribution of L1-retroelement-enriched CENP-A clusters within a functional human neocentromere. Hum Mol Genet 14:85–93.

Clarke L, Carbon J. 1985. The structure and function of yeast centromeres. Annu Rev Genet 19:29–55.

Clavijo B, Garcia Accinelli G, Wright J, Heavens D, Barr K, Yanes L, Di-Palma F. 2017. W2RAP: a pipeline for high quality, robust assemblies of large complex genomes from short read data. In: bioRxiv.

Danecek P, Bonfield JK, Liddle J, Marshall J, Ohan V, Pollard MO, Whitwham A, Keane T, McCarthy SA, Davies RM, et al. 2021. Twelve years of SAMtools and BCFtools. Gigascience 10.

Doucet AJ, Wilusz JE, Miyoshi T, Liu Y, Moran JV. 2015. A 3’ Poly(A) Tract Is Required for LINE-1 Retrotransposition. Mol Cell 60:728–741.

Dudchenko O, Batra SS, Omer AD, Nyquist SK, Hoeger M, Durand NC, Shamim MS, Machol I, Lander ES, Aiden AP, et al. 2017. De novo assembly of the <em>Aedes aegypti</em> genome using Hi-C yields chromosome-length scaffolds. Science 356:92.

Dudchenko O, Shamim S, Batra S, Durand N, Musial N, Mostofa R, Pham M, Hilaire B, Yao W, Stamenova E, et al. 2018. The Juicebox Assembly Tools module facilitates de novo assembly of mammalian genomes with chromosome-length scaffolds for under $1000.

Feng J, Liu T, Qin B, Zhang Y, Liu XS. 2012. Identifying ChIP-seq enrichment using MACS. Nat Protoc 7:1728–1740.

Fishman L, Saunders A. 2008. Centromere-associated female meiotic drive entails male fitness costs in monkeyflowers. Science 322:1559–1562.

Garagna S, Marziliano N, Zuccotti M, Searle JB, Capanna E, Redi CA. 2001. Pericentromeric organization at the fusion point of mouse Robertsonian translocation chromosomes. Proc Natl Acad Sci U S A 98:171–175.

Garagna S, Page J, Fernandez-Donoso R, Zuccotti M, Searle JB. 2014. The Robertsonian phenomenon in the house mouse: mutation, meiosis and speciation. Chromosoma 123:529–544.

Giulotto E, Raimondi E, Sullivan KF. 2017. The Unique DNA Sequences Underlying Equine Centromeres. Prog Mol Subcell Biol 56:337–354.

Ivancevic AM, Kortschak RD, Bertozzi T, Adelson DL. 2016. LINEs between Species: Evolutionary Dynamics of LINE-1 Retrotransposons across the Eukaryotic Tree of Life. Genome Biol Evol 8:3301–3322.

Iwata-Otsubo A, Dawicki-McKenna JM, Akera T, Falk SJ, Chmátal L, Yang K, Sullivan BA, Schultz RM, Lampson MA, Black BE. 2017. Expanded Satellite Repeats Amplify a Discrete CENP-A Nucleosome Assembly Site on Chromosomes that Drive in Female Meiosis. Curr Biol 27:2365-2373.e2368.

Jónsson H, Schubert M, Seguin-Orlando A, Ginolhac A, Petersen L, Fumagalli M, Albrechtsen A, Petersen B, Korneliussen TS, Vilstrup JT, et al. 2014. Speciation with gene flow in equids despite extensive chromosomal plasticity. Proc Natl Acad Sci U S A 111:18655–18660.

Kalbfleisch TS, Rice ES, DePriest MS, Walenz BP, Hestand MS, Vermeesch JR, O Connell BL, Fiddes IT, Vershinina AO, Saremi NF, et al. 2018. Improved reference genome for the domestic horse increases assembly contiguity and composition. Commun Biol 1:197.

Kasinathan S, Henikoff S. 2018. Non-B-Form DNA Is Enriched at Centromeres. Mol Biol Evol 35:949–962.

Kent WJ. 2002. BLAT--the BLAST-like alignment tool. Genome Res 12:656–664.

Klein SJ, O’Neill RJ. 2018. Transposable elements: genome innovation, chromosome diversity, and centromere conflict. Chromosome Res 26:5–23.

Kursel LE, Malik HS. 2018. The cellular mechanisms and consequences of centromere drive. Curr Opin Cell Biol 52:58–65.

Langmead B, Salzberg SL. 2012. Fast gapped-read alignment with Bowtie 2. Nat Methods 9:357–359.

Langmead B, Trapnell C, Pop M, Salzberg SL. 2009. Ultrafast and memory-efficient alignment of short DNA sequences to the human genome. Genome Biol 10:R25.

Longo MS, Carone DM, Green ED, O’Neill MJ, O’Neill RJ, Program NCS. 2009. Distinct retroelement classes define evolutionary breakpoints demarcating sites of evolutionary novelty. BMC Genomics 10:334.

Marshall OJ, Chueh AC, Wong LH, Choo KH. 2008. Neocentromeres: new insights into centromere structure, disease development, and karyotype evolution. Am J Hum Genet 82:261–282.

Musilova P, Kubickova S, Vahala J, Rubes J. 2013. Subchromosomal karyotype evolution in Equidae. Chromosome Res 21:175–187.

Myka JL, Lear TL, Houck ML, Ryder OA, Bailey E. 2003. Homologous fission event(s) implicated for chromosomal polymorphisms among five species in the genus Equus. Cytogenet Genome Res 102:217–221.

Nergadze SG, Belloni E, Piras FM, Khoriauli L, Mazzagatti A, Vella F, Bensi M, Vitelli V, Giulotto E, Raimondi E. 2014. Discovery and comparative analysis of a novel satellite, EC137, in horses and other equids. Cytogenet Genome Res 144:114–123.

Nergadze SG, Piras FM, Gamba R, Corbo M, Cerutti F, McCarter JGW, Cappelletti E, Gozzo F, Harman RM, Antczak DF, et al. 2018. Birth, evolution, and transmission of satellite-free mammalian centromeric domains. Genome Res 28:789–799.

Orlando L, Ginolhac A, Zhang G, Froese D, Albrechtsen A, Stiller M, Schubert M, Cappellini E, Petersen B, Moltke I, et al. 2013. Recalibrating Equus evolution using the genome sequence of an early Middle Pleistocene horse. Nature 499:74–78.

Phanstiel DH, Boyle AP, Araya CL, Snyder MP. 2014. Sushi.R: flexible, quantitative and integrative genomic visualizations for publication-quality multi-panel figures. Bioinformatics 30:2808–2810.

Piras FM, Nergadze SG, Magnani E, Bertoni L, Attolini C, Khoriauli L, Raimondi E, Giulotto E. 2010. Uncoupling of satellite DNA and centromeric function in the genus Equus. PLoS Genet 6:e1000845.

Piras FM, Nergadze SG, Poletto V, Cerutti F, Ryder OA, Leeb T, Raimondi E, Giulotto E. 2009. Phylogeny of horse chromosome 5q in the genus Equus and centromere repositioning. Cytogenet Genome Res 126:165–172.

Plohl M, Luchetti A, Mestrović N, Mantovani B. 2008. Satellite DNAs between selfishness and functionality: structure, genomics and evolution of tandem repeats in centromeric (hetero)chromatin. Gene 409:72–82.

Plohl M, Meštrović N, Mravinac B. 2014. Centromere identity from the DNA point of view. Chromosoma 123:313–325.

Purgato S, Belloni E, Piras FM, Zoli M, Badiale C, Cerutti F, Mazzagatti A, Perini G, Della Valle G, Nergadze SG, et al. 2015. Centromere sliding on a mammalian chromosome. Chromosoma 124:277–287.

Pérez-Wohlfeil E, Diaz-Del-Pino S, Trelles O. 2019. Ultra-fast genome comparison for large-scale genomic experiments. Sci Rep 9:10274.

Ramírez F, Ryan DP, Grüning B, Bhardwaj V, Kilpert F, Richter AS, Heyne S, Dündar F, Manke T. 2016. deepTools2: a next generation web server for deep-sequencing data analysis. Nucleic Acids Res 44:W160–165.

Ransom J, Kaczensky P. 2016. Wild equids: Ecology, management, and conservation.

Roberti A, Bensi M, Mazzagatti A, Piras FM, Nergadze SG, Giulotto E, Raimondi E. 2019. Satellite DNA at the Centromere is Dispensable for Segregation Fidelity. Genes (Basel) 10.

Rocchi M, Archidiacono N, Schempp W, Capozzi O, Stanyon R. 2012. Centromere repositioning in mammals. Heredity (Edinb) 108:59–67.

Schloss PD, Westcott SL, Ryabin T, Hall JR, Hartmann M, Hollister EB, Lesniewski RA, Oakley BB, Parks DH, Robinson CJ, et al. 2009. Introducing mothur: open-source, platform-independent, community-supported software for describing and comparing microbial communities. Appl Environ Microbiol 75:7537–7541.

Sultana T, van Essen D, Siol O, Bailly-Bechet M, Philippe C, Zine El Aabidine A, Pioger L, Nigumann P, Saccani S, Andrau JC, et al. 2019. The Landscape of L1 Retrotransposons in the Human Genome Is Shaped by Pre-insertion Sequence Biases and Post-insertion Selection. Mol Cell 74:555-570.e557.

Talbert PB, Henikoff S. 2020. What makes a centromere? Exp Cell Res 389:111895.

Trifonov VA, Musilova P, Kulemsina AI. 2012. Chromosome evolution in Perissodactyla. Cytogenet Genome Res 137:208–217.

Trifonov VA, Stanyon R, Nesterenko AI, Fu B, Perelman PL, O’Brien PC, Stone G, Rubtsova NV, Houck ML, Robinson TJ, et al. 2008. Multidirectional cross-species painting illuminates the history of karyotypic evolution in Perissodactyla. Chromosome Res 16:89–107.

Ventura M, Weigl S, Carbone L, Cardone MF, Misceo D, Teti M, D’Addabbo P, Wandall A, Björck E, de Jong PJ, et al. 2004. Recurrent sites for new centromere seeding. Genome Res 14:1696–1703.

Wade CM, Giulotto E, Sigurdsson S, Zoli M, Gnerre S, Imsland F, Lear TL, Adelson DL, Bailey E, Bellone RR, et al. 2009. Genome sequence, comparative analysis, and population genetics of the domestic horse. Science 326:865–867.

Wei KH, Reddy HM, Rathnam C, Lee J, Lin D, Ji S, Mason JM, Clark AG, Barbash DA. 2017. A Pooled Sequencing Approach Identifies a Candidate Meiotic Driver in. Genetics 206:451–465.

Yadav V, Sun S, Billmyre RB, Thimmappa BC, Shea T, Lintner R, Bakkeren G, Cuomo CA, Heitman J, Sanyal K. 2018. RNAi is a critical determinant of centromere evolution in closely related fungi. Proc Natl Acad Sci U S A 115:3108–3113.

Zhang Y, Liu T, Meyer CA, Eeckhoute J, Johnson DS, Bernstein BE, Nusbaum C, Myers RM, Brown M, Li W, et al. 2008. Model-based analysis of ChIP-Seq (MACS). Genome Biol 9:R137.

