## supplementary material for "Neocentromere formation through Robertsonian fusion and centromere repositioning during the evolution of zebras"

### Table of Contents

**Supplementary Figure S1:** Pairwise genome comparison between EquCab3.0 and Equus_quagga_HiC scaffolds

**Supplementary Figure S2:** Sequence analysis of satellite-free centromeres in Burchell's and Grevy's zebras

**Supplementary Table S1:** Burchell's (EBU) and Grevy's (EGR) zebras neocentromeres identified in EquCab3.0 and Equus_quagga_HiC assemblies

**Supplementary Table S2.** Information on the correction of the Equus_quagga_HiC assembly

**Supplementary Table S3.** Coordinates of Burchell's zebra CENP-A binding domains in Equus_quagga_cen

**Supplementary Table S4.** Information on the assembly of the chimeric reference genome obtained by inserting the centromeric *de novo* contigs of Grevy's zebra (EGR) in the Burchell's zebra (EBU) genome

**Supplementary Table S5.** Coordinates of Grevy's zebra CENP-A binding domains of EBU-EGR_cen

**Supplementary Table S6:** Sequence analysis of Robertsonian fusions in Burchell's and Grevy's zebras

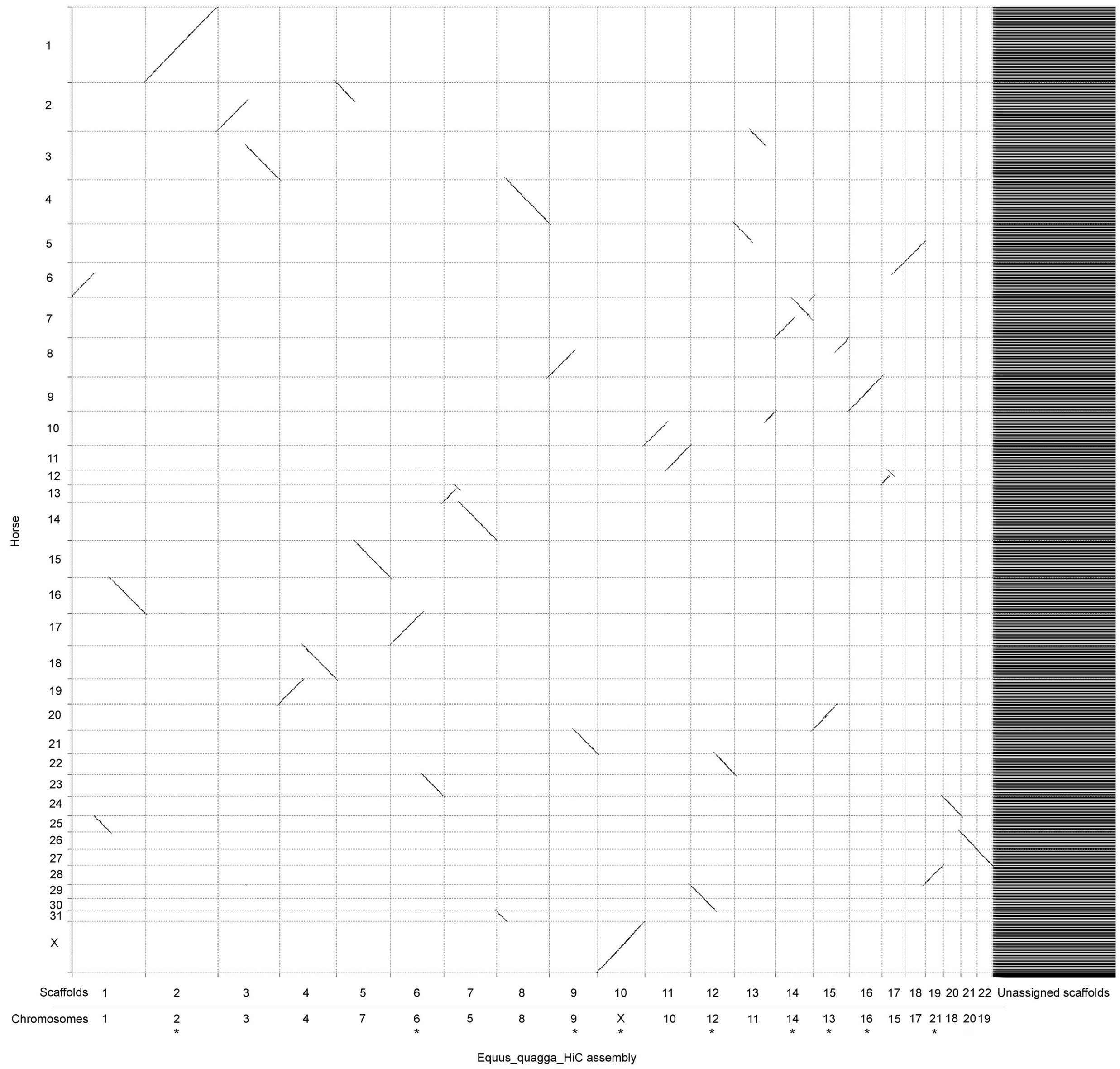

**Supplementary Figure S1. Pairwise genome comparison between EquCab3.0 and Equus_quagga_HiC scaffolds.** Aligned segments between horse chromosomes (y-axis) and Equus_quagga_HiC scaffolds (x-axis) are represented as lines. For each Burchell's zebra scaffold, the chromosome number is reported below. Asterisks indicate scaffolds with a reverse orientation compared to the direction previously determined by cytogenetic analysis (Musilova et al. 2013).

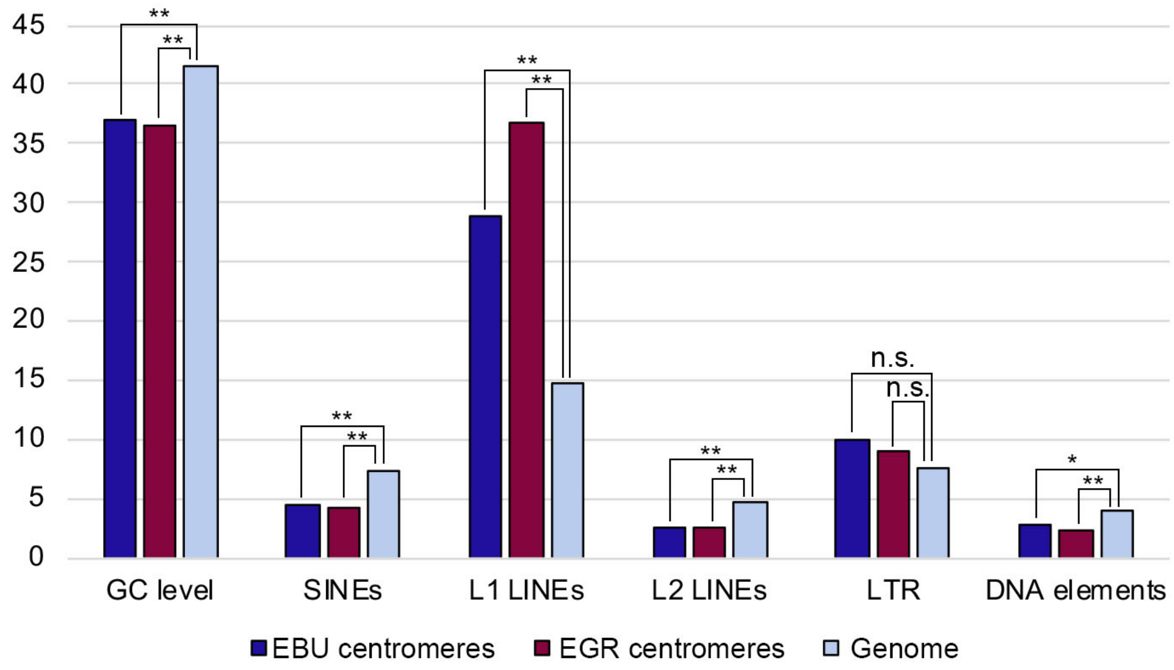

**Supplementary Figure S2. Sequence analysis of satellite-free centromeres in Burchell's and Grevy's zebras.** Sequences from the assembled satellite-free domains from Burchell's zebra and Grevy's zebras were analyzed for GC, SINEs, L1 LINEs, L2 LINEs, LTRs and DNA elements content. Centromeres containing amplicons (EBU17, EBU18, EGR1 and EGR16) were excluded from the calculations. The values obtained from each species were compared with genome wide average. Single or double asterisks indicate statistically significant differences, with a p-value < 0.05 and a p-value < 0.01 in the t-test, respectively. Differences in LTR content are not significant (n. s.).

| Chromosome with satellite-free centromere | Coordinates on EquCab3.0 | Coordinates on Equus_quagga_HiC |
|---|---|---|
| EBU4 | chr18:7,082,654-7,316,926 | HiC_scaffold_4:65,102,091-65,337,512 |
| EBU6 | unplaced | HiC_scaffold_6:77,556,520-77,775,558 |
| EBU7 | chr15:2,133,002-2,278,258 | unplaced |
| EBU8 | chr4:26,077,823-26,165,783 | unplaced |
| EBU9 | chr21:522,698-767,587 | HiC_scaffold_9:61,619,924-61,706,473 |
| EBU10 | chr10:33,057,551-33,418,701 | HiC_scaffold_11:49,436,019-49,726,883 |
| EBU12 | chr22:11,544,729-11,750,509 | HiC_scaffold_12:70,106,917-70,158,118 |
| EBU14 | chr7:53,432,077-53,821,841 | unplaced |
| EBU15 | chr12:19,972,567-20,216,155 | unplaced |
| EBU16 | chr9:34,655,231-34,911,697 | HiC_scaffold_16:46,963,937-47,211,788 |
| EBU17 | chr5:71,810,780-71,891,232 | unplaced |
| EBU18 | chr24:16,335,232-16,404,215 | unplaced |
| EBU20 | chr26:6,267,448-6,525,562 | HiC_scaffold_21:4,941,599-5,111,546 |
| EBU21 | chr28:2,659,126-3,170,658 | HiC_scaffold_19:41,292,526-41,545,076 |
| EBUX | chrX:48,174,202-48,759,430 | HiC_scaffold_10:72,225,909-72,523,907 |
| EGR1 | chr6:39,319,848-39,471,120 | unplaced |
| EGR3 | unplaced | HiC_scaffold_3:69,386,723-69,715,629 |
| EGR4 | chr7:53,638,050-53,825,901 | unplaced |
| EGR5 | chr13:7,025,768-7,733,356 | HiC_scaffold_7:33,318,974-33,837,267 |
| EGR6 | unplaced | HiC_scaffold_6:77,070,413-77,278,129 |
| EGR9 | chr18:7,290,148-7,710,732 | HiC_scaffold_4:65,289,176-65,624,658 |
| EGR10 | chr10:33,057,551-33,418,701 | HiC_scaffold_11:49,506,776-49,717,350 |
| EGR11 | chr21:431,291-949,576 | HiC_scaffold_9:61,472,997-62,035,928 |
| EGR15 | chr9:34,715,614-35,008,135 | HiC_scaffold_16:46,855,502-47,187,296 |
| EGR16 | chr24:16,322,837-16,585,836 | unplaced |
| EGR19 | chr27:630,777-1,130,366 | HiC_scaffold_22:1-50,223 |
| EGR20 | chr26:6,164,177-6,631,723 | HiC_scaffold_21:4,800,744-5,149,972 |
| EGRX | chrX:48,526,558-48,760,648 | HiC_scaffold_10:72,228,942-72,494,136 |

**Supplementary Table S1. Burchell's (EBU) and Grevy's (EGR) zebras neocentromeres identified in EquCab3.0 and Equus_quagga_HiC assemblies.**

| Chromosome | Scaffold | Direction | Region removed from Equus_quagga_HiC | | | Length of contig (bp)* |
|---|---|---|---|---|---|---|
| | | | **Start** | **End** | **Length (bp)** | |
| 2 | HiC_scaffold_2 | reverse | - | - | - | - |
| 4 | HiC_scaffold_4 | correct | 65000352 | 65401208 | 400857 | 400755 |
| 6 | HiC_scaffold_6 | reverse | 77574550 | 77804410 | 229861 | 227903 |
| 7 | HiC_scaffold_5 | correct | 46746777 | 46791095 | 44319 | 375668 |
| 8 | HiC_scaffold_8 | correct | 48901045 | 48901045 | 0 | 103276 |
| 9 | HiC_scaffold_9 | reverse | 61620303 | 61725911 | 105609 | 242915 |
| 10 | HiC_scaffold_11 | correct | 49457111 | 49768088 | 310978 | 353087 |
| 12 | HiC_scaffold_12 | reverse | 70109432 | 70155316 | 45885 | 265579 |
| 13 | HiC_scaffold_15 | reverse | - | - | - | - |
| 14 | HiC_scaffold_14 | reverse | 85528671 | 85544330 | 15660 | 348903 |
| 15 | HiC_scaffold_17 | correct | 13561248 | 13573542 | 12295 | 259470 |
| 16 | HiC_scaffold_16 | reverse | 46919921 | 47310400 | 390480 | 408827 |
| 17 | HiC_scaffold_18 | correct | 23749281 | 24047281 | 298001 | 351134 |
| 18 | HiC_scaffold_20 | correct | 14819873 | 14828451 | 8579 | 67126 |
| 20 | HiC_scaffold_21 | correct | 4738750 | 5138240 | 399491 | 443694 |
| 21 | HiC_scaffold_19 | reverse | 41323561 | 41526650 | 203090 | 202233 |
| X | HiC_scaffold_10 | reverse | 72233227 | 72521723 | 288497 | 288000 |

**Supplementary Table S2. Information on the correction of the Equus_quagga_HiC assembly.**
* Number of *de novo* sequenced nucleotides

| EBU chromosome | Coordinates of CENP-A binding domain in Equus_quagga_cen |
|---|---|
| 4 | chr4:65106754-65323307 |
| 6 | chr6:51124660-51321792 |
| 7 | chr7:46856363-47004764 |
| 8 | chr8:48900805-48987137 |
| 9 | chr9:54767829-54993986 |
| 10 | chr10:49444320-49777808 |
| 12 | chr12:35440618-35616511 |
| 14 | chr14:5429479-5706897 |
| 15 | chr15:13559837-13817437 |
| 16 | chr16:32426231-32674569 |
| 17 | chr17:23902991-23932778 |
| 18 | chr18:14827756-14854616 |
| 20 | chr20:4937529-5162913 |
| 21 | chr21:1717523-1950623 |
| X | chrX:41973637-42275187 |

**Supplementary Table S3. Coordinates of Burchell's zebra CENP-A binding domains in Equus_quagga_cen.**

| EGR chromosome | EBU chromosome | Region removed from Equus_quagga_cen | | | Length of contig (bp)* |
|---|---|---|---|---|---|
| | | Start | End | Length (bp) | |
| 1 | 1 | 44891973 | 44896035 | 4063 | 60495 |
| 3 | 3 | 69386015 | 69712952 | 326938 | 216107 |
| 4 | 14 | 5354804 | 5582851 | 228048 | 179922 |
| 5 | 5 | 33345422 | 33817503 | 472082 | 369450 |
| 6 | 6 | 51636496 | 51815413 | 178918 | 159184 |
| 9 | 4 | 65326865 | 65697798 | 370934 | 279771 |
| 10 | 10 | 49509838 | 49759385 | 249548 | 231272 |
| 11 | 9 | 54485431 | 54745834 | 260404 | 212667 |
| 15 | 16 | 32478494 | 32766972 | 288479 | 291157 |
| 16 | 18 | 14821244 | 14894826 | 73583 | 70403 |
| 19 | 19 | 1 | 50879 | 50879 | 125769 |
| 20 | 20 | 4823351 | 5182193 | 358843 | 363434 |
| X | X | 42114855 | 42271186 | 156332 | 176070 |

**Supplementary Table S4. Information on the assembly of the chimeric reference genome obtained by inserting the centromeric *de novo* contigs of Grevy's zebra (EGR) in the Burchell's zebra (EBU) genome.** * Number of *de novo* sequenced nucleotides

| EGR chromosome | Coordinates of CENP-A binding domain on EBU_EGR_cen |
|---|---|
| 1 | EBU1_EGR1cen:44,891,673-44,951,299 |
| 3 | EBU3_EGR3cen:69,381,362-69,606,129 |
| 4 | EBU14_EGR4cen:5,387,653-5,507,680 |
| 5 | EBU5_EGR5cen:33,333,014-33,708,321 |
| 6 | EBU6_EGR6cen:51,636,012-51,796,764 |
| 9 | EBU4_EGR9cen:65,311,665-65,604,244 |
| 10 | EBU10_EGR10cen:49,506,965-49,743,657 |
| 11 | EBU9_EGR11cen:54,490,698-54,700,611 |
| 15 | EBU16_EGR15cen:32,486,107-32,774,801 |
| 16 | EBU18_EGR16cen:14,825,848-14,885,596 |
| 19 | EBU19_EGR19cen:1-128,974 |
| 20 | EBU20_EGR20cen:4,816,455-5,191,599 |
| X | EBUX_EGRXcen:42,102,845-42,290,870 |

**Supplementary Table S5. Coordinates of Grevy’s zebra CENP-A binding domains of EBU_EGR_cen.**

| Robertsonian chromosome | ECA orthologies | Element with satellite-free centromere | Coordinate of the fusion region | Distance between satellite-free centromere and fusion region | Coordinates and family of satellite arrays | Cytogenetic localizations of satellite arrays |
|---|---|---|---|---|---|---|
| EBU1 | 6q/25/16 | - | chr1:55,549,823-55,561,606 | - | - | 2PI (Piras et al. 2010) |
| EGR1 | | 6q | | 10 Mb | | No signal |
| EBU3 | 2q/3q | - | chr3:68,907,682-70,420,406 | - | - | 2PI (Piras et al. 2010) |
| EGR3 | | 3q | | 450 kb | | No signal |
| EBU4 | 19/18 | 18 | chr4:58,678,012-58,683,883 | 6.5 Mb | chr4: 58,678,005-58,683,891 (2PI) | 2PI (Piras et al. 2010) |
| EGR9 | 22/18 | 18 | n.d. | n.d. | n.d. | 2PI (Piras et al. 2010) |
| EBU5 | 13/14 | - | chr5:38,512,870-38,539,229 | - | chr5:27,294,043-27,636,381 chr5: 37,874,236-38,582,889 (2PI) | 2PI (Piras et al. 2010) |
| EGR5 | | 13 | | 4.8 Mb | | No signal |
| EBU6 | 23/17 | 23 | chr6:51,032,681-52,030,540 | 500 kb | - | No signal |
| EGR6 | | | | Adjacent | | No signal |
| EBU7 | 2p/15 | 15 | chr7:44,890,018-45,000,675 | 2 Mb | chr7:44,916,147-44,956,281 (137sat; 2PI) | 137sat (Nergadze et al. 2014) |
| EGR7 | | - | | - | | 2PI (Piras et al. 2010) |
| EBU8 | 31/4 | 4 | chr8: 24,702,681-24,703,180 | 24 Mb | - | No signal |
| EBU9 | 21/8q | 21 | chr9:55,017,287-55,269,713 | adjacent | chr9:55,269,928-55,332,292 (137sat; 2PI) | 137sat (Nergadze et al. 2014) |
| EGR11 | 21/19 | | | 260 kb | | No signal |
| EBU10 | 10q/11 | 10q | chr10: 53,777,983-53,889,885 | 4 Mb | chr10: 53,715,496-54,716,757 (137sat; 2PI) | 137sat (Nergadze et al. 2014); 2PI (Piras et al. 2010) |
| EGR10 | | | | | | 137sat (Nergadze et al. 2014) |
| EBU12 | 22/30/29 | 22 | chr12:46,221,267-46,221,767 | 10 Mb | chr12:45,666,426-45,687,678 (2PI) | 2PI (Piras et al. 2010) |
| EBU15 | 12/6p | 12 | chr15:28,432,824-28,452,154 | 14 Mb | chr15:13,871,939-13,882,032 (2PI) | 2PI (Piras et al. 2010) |

**Supplementary Table S6. Sequence analysis of Robertsonian fusions in Burchell's and Grevy's zebras.**
